# Metabolomic and Proteomic Stratification of Equine Osteoarthritis

**DOI:** 10.1101/2020.05.04.077305

**Authors:** James R Anderson, Marie M Phelan, Eva Caamaño-Gutiérrez, Peter D Clegg, Luis M Rubio-Martinez, Mandy J Peffers

## Abstract

Osteoarthritis (OA) is characterised by loss of articular cartilage, synovial membrane dysfunction and subchondral sclerosis. Few studies have used a global approach to stratify equine synovial fluid (SF) molecular profiles according to OA severity. SF was collected from 58 metacarpophalangeal (MCP) and metatarsophalangeal joints of racing Thoroughbred horses (Hong Kong Jockey Club; HKJC) and 83 MCP joints of mixed breed horses from an abattoir and equine hospital (biobank). Joints were histologically and macroscopically assessed for OA severity. For proteomic analysis, native SF and SF loaded onto ProteoMiner™ equalisation columns, to deplete high abundant proteins, were analysed using liquid chromatography-tandem mass spectrometry (LC-MS/MS) and label-free quantification. Validation of selected differentially expressed proteins was undertaken using clinical SF collected during diagnostic investigations. Native SF metabolites were analysed using 1D ^1^H Nuclear Magnetic Resonance (NMR). 1,834 proteins and 40 metabolites were identified in equine SF. Afamin levels decreased with synovitis severity and four uncharacterised proteins decreased with OA severity. Gelsolin and lipoprotein binding protein decreased with OA severity and apolipoprotein A1 levels increased for mild and moderate OA. Within the biobank, glutamate levels decreased with OA severity and for the HKJC cohort, 2-aminobutyrate, alanine and creatine increased with severity. Proteomic and metabolomic integration was undertaken using linear regression via Lasso penalisation modelling, incorporating 29 variables (R^2^=0.82) with principal component 2 able to discriminate advanced OA from earlier stages, predominantly driven by H9GZQ9, F6ZR63 and alanine. Combining biobank and HKJC datasets, discriminant analysis of principal components modelling prediction was good for mild OA (90%). This study has stratified equine OA using both metabolomic and proteomic SF profiles and identified a panel of markers of interest which may be applicable to grading OA severity. This is also the first study to undertake computational integration of NMR metabolomic and LC-MS/MS proteomic datasets of any biological system.

## Introduction

The age-related degenerative musculoskeletal condition osteoarthritis (OA) is mainly characterised by articular cartilage degradation, synovitis, subchondral bone sclerosis and abnormal bone proliferation (1, 2). OA is an important welfare issue for equids, with up to 60% of lameness cases attributed to OA, leading to substantial morbidity and mortality (3–5). Although it is known that degradation of the extracellular matrix (ECM) is driven by increased enzymatic activity of multiple matrix metalloproteinases (MMPs) and a disintegrin and metalloproteinases with thrombospondin motifs (ADAMTSs), the underlying pathogenesis of OA is yet to be fully understood (6–8). Currently, equine OA is predominantly diagnosed through radiography, however due to the slow onset of the condition this often leads to substantial pathology of the joint and articular cartilage degradation prior to diagnosis (9). There is therefore a need to develop accurate biomarkers of early OA which can be applied to a clinical setting and allow for timely intervention, as well as improving our understanding of OA pathogenesis and identifying potentially novel therapeutic targets.

Currently, no equine OA-specific biomarkers have been identified to aid an early clinical diagnosis (10). For human OA diagnosis, increased synovial fluid (SF) abundances of both matrix metalloproteinases (MMPs) and cartilage oligomeric matrix protein (COMP) have been identified as markers of interest (11). For horses, MMP activity has also shown OA diagnostic potential, however studies investigating COMP levels have shown conflicting results with SF COMP abundance unable to stage equine OA (12–19). However, these markers are generated following significant joint pathology, including substantial articular cartilage degradation, and thus there is a need to identify markers at an earlier disease stage, when intervention would be most beneficial.

Palmar/plantar osteochondral disease (POD), located at the distal condyles of metacarpal III and metatarsal III, is a highly prevalent pathology of racehorses, resultant of repetitive joint overload during cyclic locomotion at high-speed (20–22). POD lesions range in severity, from mild to end-stage disease, and as the disease originates within the subchondral bone, provides an effective model to investigate subchondral bone mediated OA (23).

The inner layer of the joint capsule consists of a one-cell thick lining of synoviocytes within a hyaluronic acid and collagen matrix, called the synovial membrane, which produces SF (24, 25). SF primarily acts as a lubricant within the joint, protecting hyaline articular cartilage surfaces (26). However, SF also provides a pool of nutrients for surrounding tissues and a medium of cellular communication with the semi-permeable synovial membrane allowing passive protein transfer and synoviocytes secreting regulatory cytokines and growth factors (24–28). Due to the close relationship, both in terms of location and biological communication that SF holds with surrounding tissues which are primarily altered during OA, and its accessibility, SF can provide a unique source of chemical information and holds great promise for biomarker discovery (10, 29).

Various studies have utilised ^1^H nuclear magnetic resonance (NMR) to investigate metabolite markers associated with OA in various species, including pigs, dogs, humans and horses (10, 30–35). Lacitignola *et al*. identified a panel of ten metabolites which were elevated in equine OA SF compared to a normal control group. This panel included alanine, acetate, N-acetyl glucosamine, pyruvate, citrate, creatine/creatinine, choline, glycerol, lactate and α-glucose. However, no studies to date have used ^1^H NMR to stratify OA to identify changes to the metabolite profile at different stages of OA severity.

Several studies have used mass spectrometry (MS) based methodologies to investigate equine OA SF (36–40). The relatively recent development of liquid chromatography tandem mass spectrometry (LC-MS/MS) has enabled a method which can quickly, with high sensitivity, quantify proteins present within biological fluids of high complexity (25). Protein biomarker discovery is however hindered by the large protein concentration dynamic range exhibited by SF, meaning that highly abundant proteins can mask those of less abundance (41, 42). ProteoMiner™ protein enrichment columns (Bio-Rad Laboratories Ltd., Hemel Hempstead, UK) have however been developed which utilise combinational ligand library technology, depleting highly abundant proteins and subsequently enriching low abundance proteins to reduce this dynamic range (43–45). This methodology has successfully been used to investigate OA in equine SF (38). However, this technique has not yet been utilised with SF to enable OA stratification, analysing SF at different OA severities.

Increased activity of MMPs, ADAMTSs, cathepsins and serine proteases during OA leads to cartilage breakdown and the generation of OA-specific peptide degradation products (neopeptides) (46–48). Numerous studies investigating equine tissue have identified potential OA neopeptides of interest using *ex-vivo* cartilage and SF (37, 38, 46, 49). As OA neopeptides have also been shown to be a driver of OA pain, identification and quantification of neopeptides has the potential to stratify OA, providing markers of early OA pathology and providing potential novel OA therapeutic and analgesic targets (50).

Within this study we have interrogated equine SF from horses with naturally occurring OA and those exhibiting POD pathology, as a subchondral bone mediated OA model using both ^1^H NMR metabolomics and LC-MS/MS proteomics to stratify equine OA. This is the first study to use both techniques on the same samples and the first to statistically integrate NMR-led metabolomics with MS-based proteomics.

### Experimental Procedures

#### Experimental Design and Statistical Rationale

A total of 141 separate equine SF samples were analysed during this study. Depending on the analysis undertaken, biological replicates ranged from n=8-34 per group for OA grading and n=17-38 per group for synovitis grading (Table 1). Joints with a microscopic or macroscopic OA grade of 0 were considered the control group, ranging from n=8-17. Metabolite and protein abundances were analysed by t-test or ANOVA, depending on the number of groups, with p values corrected for multiple testing using the Benjamini-Hochberg false discovery rate method. For NMR metabolomics and LC-MS/MS proteomics dataset integration, the Mahalanobis distance on principal components was calculated and Chi-squared testing undertaken to identify potential outliers. A linear regression, using Lasso (least absolute shrinkage and selection operator) penalisation to select the most important variables, was applied to combined datasets for both the biobank and Hong Kong Jockey Club (HKJC) sample groups. Additionally, correlations of all variables (proteins and metabolites) with macroscopic OA score were calculated, with significant variables (p < 0.05) included for further analysis. Significant variables across all analyses were then selected in a dataset, combining both biobank and HKJC results, and discriminant analysis of principal components (DAPC) was used to predict different levels of disease severity.

**Table 1.** Stratification groups according to microscopic osteoarthritis, macroscopic osteoarthritis and synovitis pathology used for synovial fluid LC-MS/MS proteomic and NMR metabolomic analysis.

Biobank
| Metabolomics | OARSI Microscopic OA Group |  |  | OARSI Macroscopic OA Group |  |  |  |
| --- | --- | --- | --- | --- | --- | --- | --- |
|  | 0 | 1 | 2 | 0 | 1 | 2 | 3 |
| Number of Donors | 8 | 34 | 28 | 14 | 27 | 21 | 14 |
| Joint | 8 x MCP | 34 x MCP | 28 x MCP | 14 x MCP | 27 x MCP | 21 x MCP | 14 x MCP |
| Mean Age (years) | 8 | 15 | 17 | 9 | 15 | 16 | 19 |
| Sex (M/F) | 1 F, 7 Unknown | 6 M, 10 F, 18 Unknown | 12 M, 10 F, 6 Unknown | 2 M, 5 F, 7 Unknown | 7 M, 12 F, 8 Unknown | 9 M, 5 F, 7 Unknown | 6 M, 3 F, 5 Unknown |

| Proteomics | OARSI Microscopic OA Group |  |  | OARSI Macroscopic OA Group |  |  |  |
| --- | --- | --- | --- | --- | --- | --- | --- |
|  | 0 | 1 | 2 | 0 | 1 | 2 | 3 |
| Number of Donors | 8 | 34 | 28 | 11 | 25 | 18 | 12 |
| Joint | 8 x MCP | 34 x MCP | 28 x MCP | 11 x MCP | 25 x MCP | 18 x MCP | 12 x MCP |
| Mean Age (years) | 8 | 15 | 17 | 9 | 15 | 16 | 20 |
| Sex (M/F) | 1 F, 7 Unknown | 6 M, 10 F, 18 Unknown | 12 M, 10 F, 6 Unknown | 1 M, 3 F, 7 Unknown | 6 M, 11 F, 8 Unknown | 6 M, 5 F, 7 Unknown | 5 M, 2 F, 5 Unknown |
| Mean Protein Concentration (mg/ml) | 5 | 5 | 4 | 5 | 4 | 5 | 6 |

Hong Kong Jockey Club
| Metabolomics | Microscopic OA Group |  |  | Macroscopic OA Group |  |  | Synovitis Group |  |  |
| --- | --- | --- | --- | --- | --- | --- | --- | --- | --- |
|  | 0 | 1 | 2 | 0 | 1 | 2 | 0 | 1 | 2 |
| Number of Donors | 17 | 20 | 13 | 13 | 25 | 12 | 1 | 38 | 17 |
| Joint | 10 x MCP, 7 x MTP | 8 x MCP, 12 x MTP | 9 x MCP, 4 x MTP | 5 x MCP, 8 x MTP | 13 x MCP, 12 x MTP | 11 x MCP, 11 x MTP | 1 x MTP | 22 x MCP, 16 x MTP | 10 x MCP, 7 x MTP |
| Mean Age (years) | 6 | 7 | 7 | 5 | 7 | 7 | 6 | 7 | 7 |
| Sex (M/F) | UNKNOWN |  |  | UNKNOWN |  |  | UNKNOWN |  |  |

| Proteomics | Microscopic OA Group |  |  | Macroscopic OA Group |  |  | Synovitis Group |  |  |
| --- | --- | --- | --- | --- | --- | --- | --- | --- | --- |
|  | 0 | 1 | 2 | 0 | 1 | 2 | 0 | 1 | 2 |
| Number of Donors | 16 | 20 | 13 | 13 | 24 | 12 | 1 | 34 | 19 |
| Joint | 9 x MCP, 7 x MTP | 8 x MCP, 12 x MTP | 9 x MCP, 4 x MTP | 5 x MCP, 8 x MTP | 12 x MCP, 12 x MTP | 11 x MCP, 11 x MTP | 1 x MTP | 18 x MCP, 16 x MTP | 10 x MCP, 9 x MTP |
| Mean Age (years) | 6 | 7 | 7 | 5 | 7 | 7 | 6 | 7 | 7 |
| Sex (M/F) | UNKNOWN |  |  | UNKNOWN |  |  | UNKNOWN |  |  |
| Mean Protein Concentration (mg/ml) | 10 | 9 | 13 | 7 | 9 | 17 | 16 | 9 | 12 |

Abbreviations: OA = Osteoarthritis; Joints, MCP = Metacarpophalangeal joint, MTP = Metatarsophalangeal joint; Sex, M = male, F = Female.

#### Sample Collection and Processing

Equine post mortem samples were collected from a commercial abattoir, The Philip Leverhulme Equine Hospital, University of Liverpool and The Hong Kong Jockey Club (HKJC) Equine Hospital. Abattoir and Philip Leverhulme Equine Hospital samples were collected from horses of mixed breed and sex and represented naturally occurring OA disease. HKJC samples were exclusively from Thoroughbred racehorses (aged 3-10 years old) with a high prevalence of POD, providing a model for subchondral bone mediated OA.

#### University of Liverpool Biobank

Following euthanasia at the abattoir (F Drury & Sons, Swindon, UK) distal equine forelimbs were transported to the University of Liverpool. Horses euthanised at the Philip Leverhulme Equine Hospital, University of Liverpool, were processed on site. Within 8 hr of euthanasia the metacarpophalangeal (MCP) joint was opened aseptically and the distal metacarpal III photographed. SF was removed using a 10 ml syringe, transferred to a plain eppendorf, centrifuged at 2,540*g* and 4°C for 5 min, supernatant removed, snap frozen in liquid nitrogen and stored at -80°C (51) (Figure S1). SF was separated into separate 1 ml eppendorfs prior to snap freezing to optimise study design prior to multi ‘omics’ analysis and integration (52). Wedge sections of articular cartilage/subchondral bone (measuring 4.0 cm x 1.5 cm x 0.5 cm) were also sampled from the distal condylar region of metacarpal III, lateral to the sagittal ridge, and placed into 4% paraformaldehyde (PFA, Sigma-Aldrich, Gillingham, UK) in phosphate buffered saline (PBS, Sigma-Aldrich) (Figure S2). Following decalcification in ethylenediaminetetraacetic acid (EDTA, Sigma-Aldrich), these were then sectioned and stained with Haematoxylin and eosin (H & E) (TCS Biosciences Ltd, Buckingham, UK) and Safranin O (BDH Chemicals, Poole, UK).

As biobank samples were obtained from a commercial abattoir and equine hospital, horses sampled were of mixed breed and sex and represented naturally occurring OA disease.

#### Hong Kong Jockey Club

HKJC samples were collected using the same collection protocols as used for the University of Liverpool equine biobank. In addition, synovial membranes were also dissected, fixed in 4% PFA and processed to prepare H & E stained histology slides. For HKJC samples, both MCP and metatarsophalangeal (MTP) joints were sampled. Following snap freezing with liquid nitrogen, frozen SF samples were shipped to Liverpool on dry ice.

#### Clinical Synovial Fluid Collection for ELISA

During clinical diagnostic investigations of horses presenting to The Philip Leverhulme Equine Hospital, University of Liverpool between 2014 and 2017, excess aspirated SF from live horses was collected and subsequently processed using the same processing steps as used for biobank and HKJC SF samples. Clinical OA was diagnosed via clinical examination, radiography and/or arthroscopy. Healthy/control SF samples were collected during clinical examinations, including diagnostic analgesia, whereby the joint involved was not identified as the cause of the current lameness. However, underlying joint pathology, not leading to a clinical manifestation, could not be ruled out. These clinical SF samples were used for enzyme-linked immunosorbent assay (ELISA) validations of altered protein abundances.

#### Osteoarthritis Pathology: Macroscopic and Microscopic Scoring

Biobank distal metacarpal III articular surfaces were assessed for macroscopic OA pathology using the equine OARSI scoring scale (53). Histological sections were also assessed for microscopic OA related pathology using the microscopic aspect of the equine OARSI scoring scale. Within the HKJC group, distal metacarpal III samples were assessed macroscopically and microscopically for OA related and POD pathology using separate published scoring scales (21, 54). Marginal remodelling and dorsal impact injury categories were excluded from the macroscopic scoring scale as these could not be scored using the joint photographs which were provided. Synovial membrane histological sections were also scored according to synovitis severity (55).

### Metabolomics

#### NMR Sample Preparation

SF was thawed over ice and centrifuged for 15 min at 13,000*g* and 4°C. 150 μl of supernatant was then diluted to produce a final volume containing 50% (v/v) SF, 40% (v/v) dd ^1^H_2_O, 100 mM PO_4_^3-^ pH 7.4 buffer (Na_2_HPO_4_, VWR International Ltd., Radnor, Pennsylvania, USA and NaH_2_PO_4_, Sigma-Aldrich) in deuterium oxide (^2^H_2_O, Sigma-Aldrich) and 0.0025% (v/v) sodium azide (NaN_3_, Sigma-Aldrich). Prepared samples were then vortexed for 1 min, centrifuged for 2 min at 13,000*g* and 4°C and, using a glass pipette, 195 μl transferred into 3 mm outer diameter NMR tubes.

#### NMR Acquisition

All SF samples were individually analysed. A 700 MHz NMR Bruker Avance III HD spectrometer with associated TCI cryoprobe and chilled Sample-Jet autosampler was used to acquire all spectra. 1D ^1^H NMR spectra, using a Carr-Purcell-Meiboom-Gill (CPMG) filter to attenuate macromolecule (e.g. protein) signals, were acquired using a standard cpmgpr1d vendor pulse sequence. Spectral acquisition was carried out at 37°C with a 4 s interscan delay, 32 transients and a 15 ppm spectral width. Software programmes Topsin 3.1 and IconNMR 4.6.7 were used for acquisition and processing, carrying out automated phasing, baseline correction and a standard vendor processing routine (exponential window function with 0.3 Hz line broadening).

#### Metabolite Annotation and Identification

All 1D ^1^H NMR spectra were scrutinised to make sure that the minimum reporting standards were met, as outlined by the Metabolomics Society (56). These quality control criteria included flat baseline correction, water suppression, and consistent line widths. Spectra which did not meet these minimum requirements were removed from all subsequent analyses. Spectra were aligned to a single formate peak at 8.46 ppm. Chenomx NMR Suite 8.2 (330-mammalian metabolite library) software was used to carry out metabolite annotations and relative abundances. Metabolite identifications were confirmed where possible using in-house 1D ^1^H NMR metabolite spectral library standards.

### Proteomics

#### Synovial Fluid Processing and Protein Assay

SF was thawed on ice and centrifuged at 4°C at 14,000*g* for 10 min. The supernatant was treated with 1μg/ml of hyaluronidase (bovine origin, Sigma-Aldrich) at 37°C for 1 hr, centrifuged at 1,000*g* for 5 min, supernatant removed and 1 ml centrifuged through a polypropylene microcentrifuge tube filter with 0.22 μm pore cellulose acetate membrane (Costar Spin-X, Corning, Tokyo, Japan) at 5,000*g* for 15 min to remove remaining insoluble particulates (29). A Pierce^®^ 660 nm protein assay (Thermo Scientific, Waltham, Massachusetts, USA) was used to determine SF protein concentrations.

#### ProteoMiner™ Column Processing

2 mg of protein was loaded onto ProteoMiner™ Small Capacity bead columns (Bio-Rad Laboratories Ltd., Hemel Hempstead, UK) to achieve peptide-based depletion of the most abundant proteins. SF samples were rotated for 2 hr at room temperature and centrifuged at 1,000*g* for 60 s. ProteoMiner™beads were then washed in PBS, rotated for 5 min and centrifuged at 1,000*g* for 60 s. The wash step was repeated two further times.

#### Protein Digestion

To assess both high and low abundance synovial proteins, both native and ProteoMiner™ processed SF were analysed. For native SF, 100 μg of protein was used for each protein trypsin digestion. 25 mM ammonium bicarbonate (Fluka Chemicals Ltd., Gillingham, UK) containing 0.05% (w/v) RapiGest (Waters, Elstree, Hertfordshire, UK) was added to both native SF and peptide bound ProteoMiner™beads to produce a final volume of 160 μl and heated at 80°C for 10 min. DL-Dithiothreitol (Sigma-Aldrich) was added (3 mM final concentration) and incubated at 60°C for 10 min. Iodoacetamide (Sigma-Aldrich) was added (9 mM final concentration) and incubated at room temperature for 30 min in the dark.

ProteoMiner™ processed samples then underwent an additional step which entailed the addition of 2 μg of Lys-C endopeptidase (FUJIFILM Wako Pure Chemical, Osaka, Japan) and incubation at 37°C for 4 hr (51). 2 μg of proteomics grade trypsin (Sigma-Aldrich) was added to all samples and rotated for 16 hr at 37°C, followed by a second trypsin supplementation for 2 hr incubation. Digests were centrifuged at 1,000*g* for 1 min, supernatant removed, trifluoroacetic acid (TFA, Sigma-Aldrich) added (0.5% (v/v) final concentration) and rotated for 30 min at 37°C. Digests were then centrifuged at 13,000*g* for 15 min at 4°C and the supernatant removed and stored at -80°C. To ensure complete protein digestion, 5 μl of each digest was analysed by 1D SDS PAGE and stained with either Coomassie Blue (Bio-Rad) or silver stain (Thermo Scientific) following manufacturer instructions (data not shown).

#### Sample Processing for Neopeptide Analysis

After 4 hr of the 16 hr tryptic digestion of ProteoMiner™ processed samples, 10 μl was removed and supplemented with TFA and stored at -80°C using the same protocol as mentioned above.

#### Label Free LC-MS/MS

All SF digest samples underwent randomisation and were individually analysed using LC-MS/MS on an UltiMate 3000 Nano LC System (Dionex/Thermo Scientific) coupled to a Q Exactive™ Quadrupole-Orbitrap instrument (Thermo Scientific). Full LC-MS/MS instrument methods are described in the supplemental data. Tryptic peptides, which were equivalent to 200 ng of protein, were loaded onto the column and run over 60 min, 90 min and 120 min LC gradients for 4 hr Lys-C + 4 hr tryptic digest ProteoMiner™ samples, native SF and 4 hr Lys-C + 16 hr + 2 hr tryptic digest ProteoMiner™ processed samples respectively. Representative ion chromatograms are shown in Figure S3.

#### LC-MS/MS Spectra Processing and Protein Identification

Progenesis™ QI 2.0 (Nonlinear Dynamics, Waters) software was used to process raw spectral files and undertake spectral alignment, peak picking, total protein abundance normalisation and peptide/protein quantification. The top ten spectra for each feature were then exported with peptide and protein identifications carried out with PEAKS^®^ Studio 8.0 (Bioinformatics Solutions Inc., Waterloo, Ontario, Canada) using the reviewed *Equus caballus* database (downloaded 29^th^ July 2016, 22,694 sequences). Search parameters included: precursor mass error tolerance, 10.0 ppm; fragment mass error tolerance, 0.01 Da; precursor mass search type, monoisotopic; enzyme, trypsin; maximum missed cleavages, 1; non-specific cleavage, none; fixed modifications, carbamidomethylation; variable modifications, oxidation or hydroxylation and oxidation (methionine). The false discovery rate (FDR) was set to 1% with only proteins identified with > 2 unique peptides used for quantitation.

#### Semi-Tryptic Peptide Identification

A ‘semi-tryptic’ search was undertaken to identify potential synovial neopeptides. PEAKS^®^ search parameters were kept the same as those used for protein identifications, apart from ‘Non-specific Cleavage’ which was changed from ‘none’ to ‘one’. The exported ‘peptide ion measurements’ file from Progenesis™ was then analysed using an online neopeptide analyser software tool to identify only semi-tryptic peptides and perform normalisation (57).

#### Batch Corrections

ProteoMiner™ processed samples for both the biobank and HKJC cohorts were run in three separate batches on the Q Exactive™ for protein analysis. To eliminate batch effects on the final analysis, a COMBAT batch correction was applied (Figure S4) (58). Metabolomic spectra were also acquired over 3 batches for both biobank and HKJC cohorts and data also underwent COMBAT batch correction.

#### ELISA Protein Validations

Differentially expressed proteins apolipoprotein A1 (ApoA1), gelsolin and lipopolysaccharide binding protein (LBP) were selected for ELISA as commercially available kits were compatible with equine samples. Equine ApoA1 (MBS034194, MyBioSource Inc., San Diego, California, USA) and equine LBP (MBS062216, MyBioSource) utilised sandwich ELISA technology using undiluted SF whilst equine gelsolin (CSB-EL009965HO, Cusabio Technology LLC, Houston, Texas, USA) was a competitive inhibition assay with SF diluted 1:1250. 3-6 dependent (and independent where stated) SF samples were analysed per group. Aliquots of 100 μl were analysed in duplicate for each sample with absorbance measured at 450 nm and protein concentrations calculated from assay standard curves. Protein concentrations were normalised to total protein.

#### Separate Dataset Statistical Analysis

SF metabolite abundances underwent median normalisation and protein abundances were normalised to the total ion current (TIC). Before multivariate analysis, both metabolite and protein datasets underwent Pareto scaling (59). All principal component analysis (PCA), ANOVA and t-tests of metabolite, protein and neopeptide abundances were conducted using MetaboAnalyst 4.0 (http://www.metaboanalyst.ca). ELISA t-tests and ANOVAs were performed using Minitab version 17. For protein analysis an additional filter of > 2 fold abundance was implemented. A p value of < 0.05 was considered statistically significant following correction for multiple testing using the Benjamini-Hochberg false discovery rate method (60). All box plots were produced using SPSS 24.

#### NMR metabolomics and LC-MS/MS Proteomics Integration

Metabolomics and proteomics datasets were initially integrated and analysed separately for the biobank and HKJC, then finally all datasets integrated to produce an overall model.

Only datasets for horses which had both metabolite and protein abundance values were used for integration. Proteomics datasets were again normalised to the TIC whilst NMR datasets were normalised via probabilistic quotient normalisation (PQN) (61). When combining ProteoMiner™ processed SF and native SF protein abundances for the same SF sample, in which the same protein had been identified in both datasets, the higher abundance was included for analysis. The Mahalanobis distance on principal components was calculated and Chi-squared testing undertaken to identify potential outliers. When combining metabolite and protein variables, categorisation was carried out in accordance to macroscopic OA scoring. A linear regression, using Lasso penalisation to select the most important variables, was applied to combined datasets for both the biobank and HKJC sample groups. Additionally, correlations of all variables (proteins and metabolites) with macroscopic OA score were calculated using the Spearman coefficient, with significant variables after correcting for False Discovery Rate (Adjusted p value < 0.05) included for further analysis. Significant variables across all analyses were then applied to a collective dataset, combing both biobank and HKJC results. This dataset was stratified into healthy, mild OA and severe OA and a DAPC model was created to predict the disease severity. The number of parameters of the model were chosen after a cross-validation process calculated using the function xvalDapc and splitting the data into train/test subsets at a 80%/20% proportion respectively. All integration analyses were undertaken using standard analytical routines within the software R and packages adegenet, ggplot2 and reshape (62–65).

#### Uncharacterised Proteins

Within this study, proteins which were considered uncharacterised were also analysed using BLAST (Basic local alignment search tool) to assess similarity of their amino acid sequence to characterised proteins within the *Equus caballus* database as well as other species (66). Search parameters included: Matrix, blosum62; threshold, 0.1 E; filtering, none; gapped, true.

#### Pathway Analysis

Pathway analysis was conducted on proteins and metabolites which were considered significant variables for DAPC when carrying out NMR metabolomic and LC-MS/MS proteomic dataset integration. Metabolite pathway analysis was conducted using the online tool KEGG for *Equus caballus* with protein pathways also analysed using KEGG, via the STRING database (67, 68). A filter of a minimum of two metabolites or proteins was set for inclusion of the relevant biological pathway.

## Results

### Stratification Groups

Histological microscopic OA scores vs macroscopic scores for biobank and HKJC samples both showed a weak positive correlation with R^2^ coefficient values of 0.12 and 0.17 respectively (Figure S5). Joints were assigned into severity groups according to microscopic OA pathology, macroscopic OA pathology and synovitis scores (Table 1). Although scored using different scoring systems, overall the HKJC samples exhibited a more severe OA phenotype. Macroscopically, scores were on average 30% of the highest possible score, compared to 14% within the biobank and microscopic scores 31% compared to 15% within the biobank. All scoring is shown in Tables S6-S10. Overall, the HKJC dataset was a younger cohort than the biobank cohort, with average ages of 6.6 ± 1.9 years and 14.3 ± 7.6 years respectively (Table S1).

### NMR Metabolomics

Overall, both the biobank and HKJC SF 1D ^1^H NMR spectra produced similar profiles, although quantile plots revealed the biobank group exhibited more variation between samples (Figure S6). In total, 40 metabolites were identified within equine SF (Table 2). For the biobank cohort, unsupervised multivariate PCA did not identify separation between OA grades based on microscopic or macroscopic scoring (Figure 1). However, when stratified according to macroscopic grade, glutamate levels were found to be differentially abundant, with lower levels identified at grades 2 and 3 compared to 1. PCA also did not identify clear separation between severity grades of OA or synovitis for HKJC SF samples (Figure 2). When stratified according to macroscopic OA pathology, three metabolites were found to be differentially abundant. 2-aminobutyrate levels were increased at grades 1 and 2 compared to 0 and alanine and creatine levels were increased with OA severity.

**Table 2.** Synovial fluid metabolites identified using Chenomx software. Metabolites which had also undergone identification using a 1D ^1^H NMR in-house spectral library were assigned to Metabolomics Standards Initiative (MSI) level 1 (129).

| Database Identifier | Metabolite Identification | Reliability |
| --- | --- | --- |
| HMDB00650 | 2-Aminobutyrate | MS Level 2 |
| HMDB00357 | 3-Hydroxybutyrate | MS Level 2 |
| HMDB00754 | 3-Hydroxyisovalerate | MS Level 2 |
| HMDB01149 | 5-Aminolevulinate | MS Level 2 |
| HMDB00042 | Acetate | MS Level 1 |
| HMDB00194 | Anserine | MS Level 2 |
| HMDB00043 | Betaine | MS Level 1 |
| HMDB00097 | Choline | MS Level 1 |
| HMDB00094 | Citrate | MS Level 1 |
| HMDB00064 | Creatine | MS Level 1 |
| HMDB01511 | Creatine phosphate | MS Level 2 |
| HMDB00562 | Creatinine | MS Level 1 |
| HMDB00122 | D-Glucose | MS Level 1 |
| HMDB04983 | Dimethyl sulfone | MS Level 2 |
| HMDB00142 | Formate | MS Level 2 |
| HMDB00123 | Glycine | MS Level 1 |
| HMDB00128 | Guanidoacetate | MS Level 2 |
| HMDB00172 | Isoleucine | MS Level 1 |
| HMDB00190 | Lactate | MS Level 1 |
| HMDB00161 | L-Alanine | MS Level 1 |
| HMDB00062 | L-Carnitine | MS Level 2 |
| HMDB00174 | L-Fucose | MS Level 2 |
| HMDB00148 | L-Glutamate | MS Level 1 |
| HMDB00641 | L-Glutamine | MS Level 1 |
| HMDB00177 | L-Histidine | MS Level 1 |
| HMDB00687 | L-Leucine | MS Level 1 |
| HMDB00159 | L-Phenylalanine | MS Level 1 |
| HMDB00158 | L-Tyrosine | MS Level 1 |
| HMDB00883 | L-Valine | MS Level 1 |
| HMDB00691 | Malonate | MS Level 2 |
| HMDB01844 | Methylsuccinate | MS Level 2 |
| HMDB31419 | N-Nitrosodimethylamine | MS Level 2 |
| HMDB00895 | O-Acetylcholine | MS Level 2 |
| HMDB00243 | Pyruvate | MS Level 1 |
| HMDB00254 | Succinate | MS Level 1 |
| HMDB00294 | Urea | MS Level 1 |
| HMDB00296 | Uridine | MS Level 1 |
| HMDB00292 | Xanthine | MS Level 2 |
| HMDB000001 | π-Methylhistidine | MS Level 2 |
| HMDB000001 | τ-Methylhistidine | MS Level 2 |

*Metabolomics Standards Initiative definitions: MS Level 1 = Identified metabolite using two or more orthogonal properties of an authentic chemical standard analysed in the researcher's laboratory. MS Level 2 = Putatively annotated metabolite which does not require matching to data for authentic chemical standards acquired within the same laboratory.*

**Figure 1.**
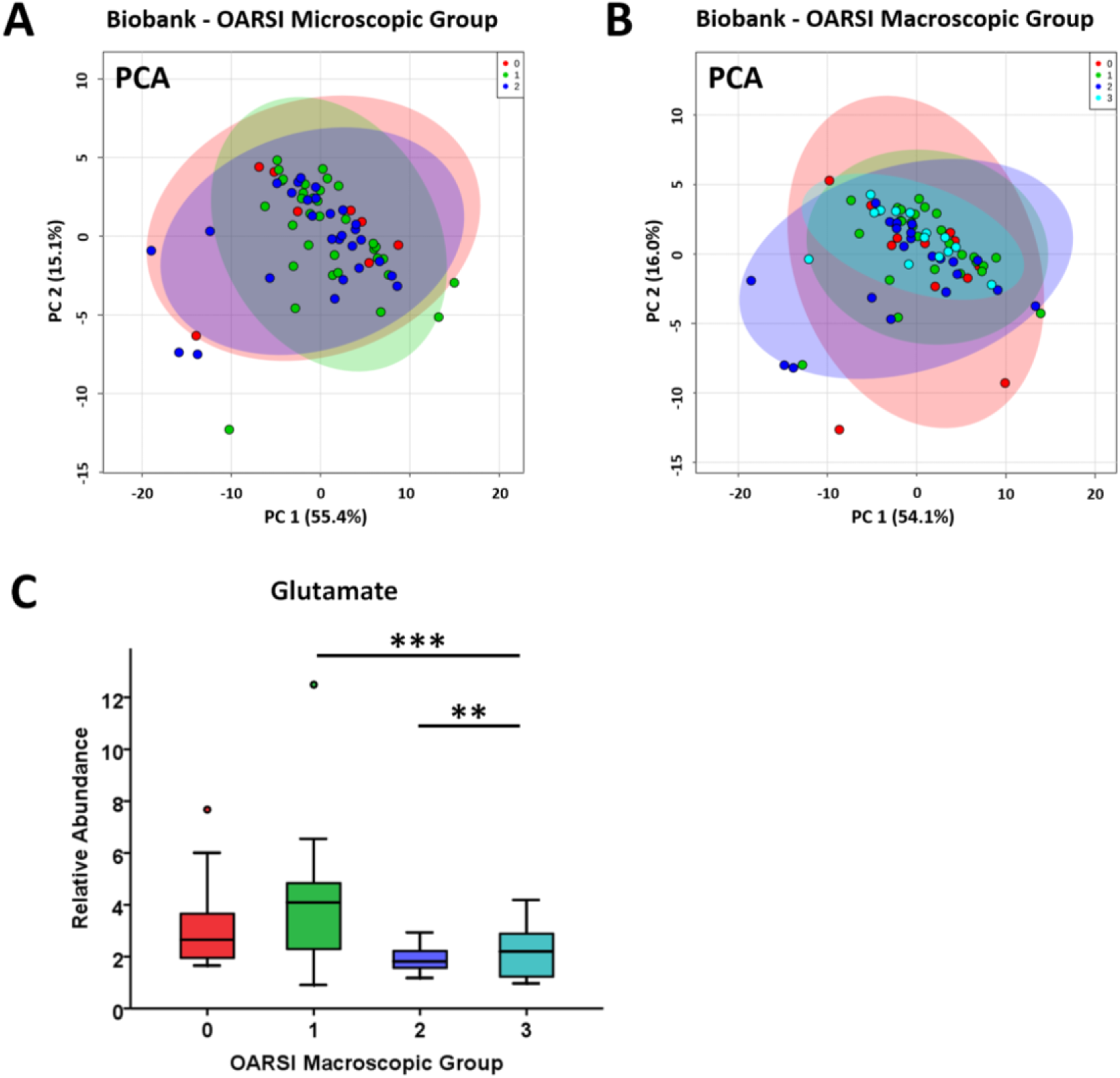
Principal component analysis (PCA) of metabolite profiles of equine biobank synovial fluid grouped according to (A) OARSI microscopic (n=70) and (B) OARSI macroscopic (n=76) grading. (C) Glutamate abundances according to macroscopic osteoarthritis grading. ANOVA: ** = p < 0.01 and *** = p < 0.001.

**Figure 2.**
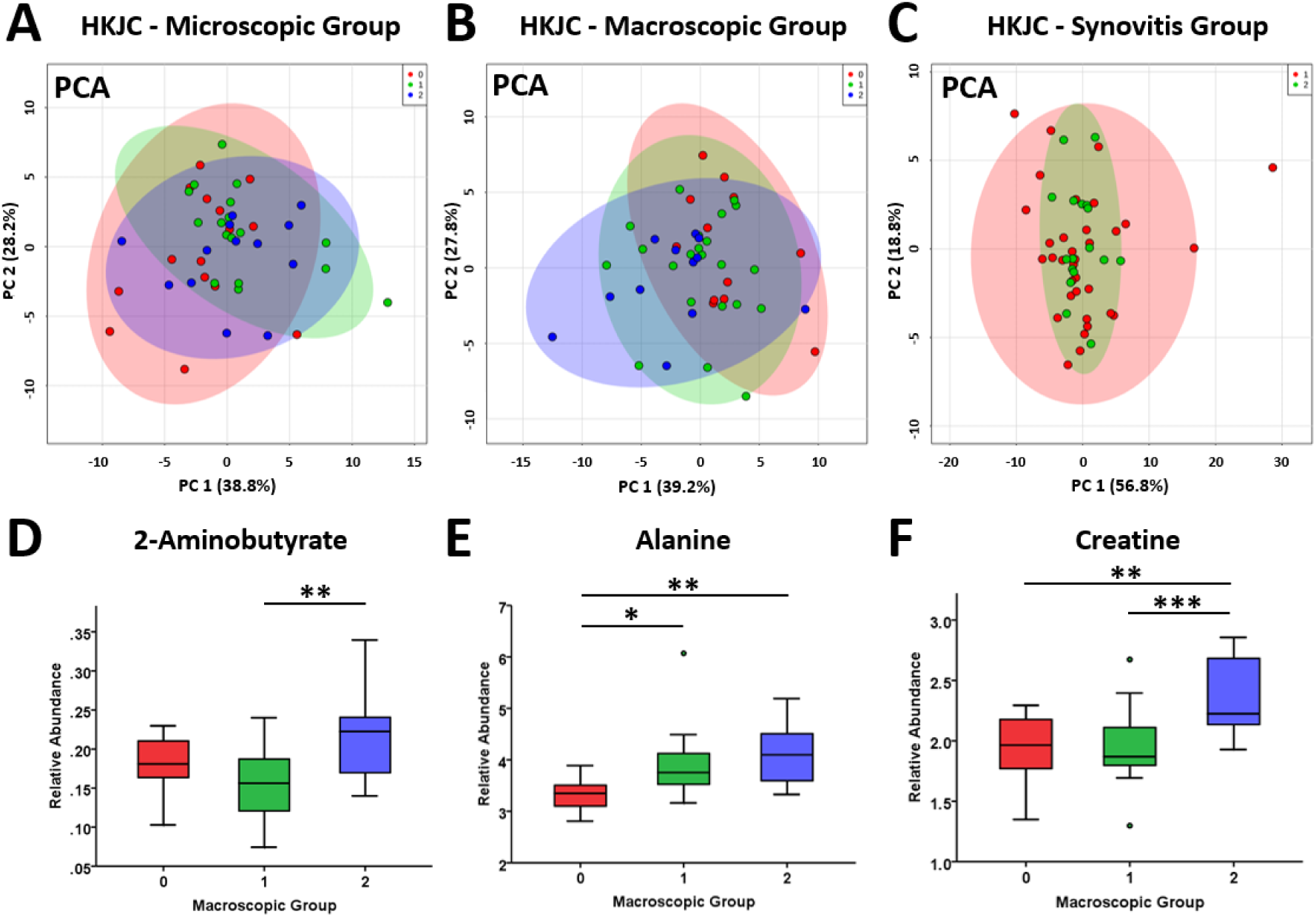
Principal component analysis (PCA) of metabolite profiles of equine Hong Kong Jockey Club (HKJC) synovial fluid (SF) grouped according to (A) OARSI microscopic (n=50), (B) OARSI macroscopic (n=50) and (C) synovitis (n=56) grading. For OARSI macroscopic grading, (D) 2-aminobutyrate, (E) alanine and (F) creatine were identified as being differentially abundant between osteoarthritis severity grades. ANOVA: * = p < 0.05, ** = p < 0.01 and *** = p < 0.001.

#### LC-MS/MS Proteomics

Within the biobank cohort, 74 native SF samples were analysed with 68 of these additionally processed using ProteoMiner™ columns. For the HKJC group, 56 native SF samples were analysed with 55 of these additionally having also undergone ProteoMiner™ processing. In total, across all samples, 1,834 proteins were identified (Figure 3). A combination of an increase in LC gradient length and ProteoMiner™ processing resulted in a 168% increase in the overall number of identified proteins compared to native SF analysis.

**Figure 3.**
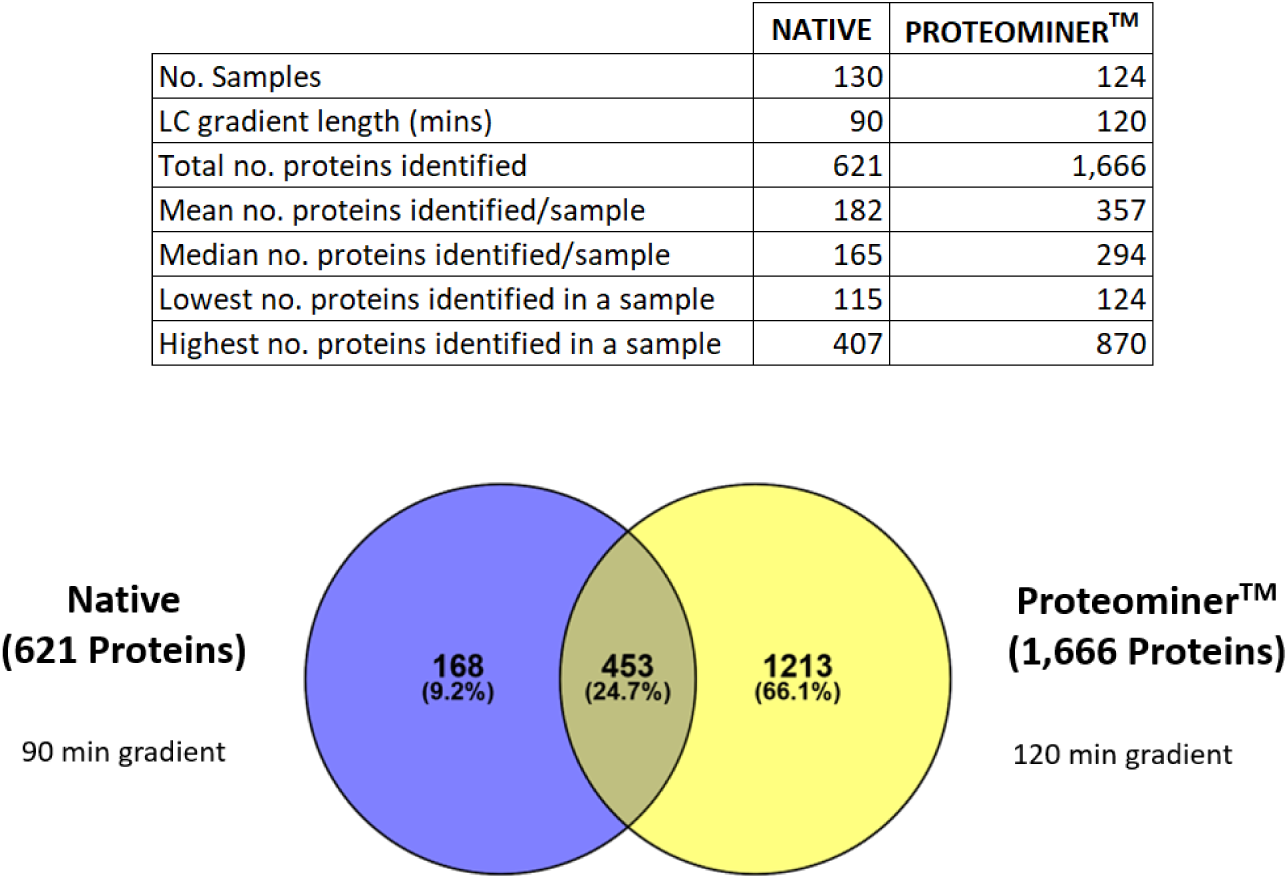
Number of proteins identified within native and ProteoMiner™ processed equine synovial fluid using liquid chromatography-tandem mass spectrometry. In total, 1,834 different proteins were identified. Search criteria included > 2 unique peptides with an FDR of 1%. LC = liquid chromatography.

Following PCA analyses, HKJC native SF samples 95 and 122 were identified as outliers and removed from further analyses. For biobank and HKJC groups, when categorised according to macroscopic OA severity, PCA identified that increased OA severity resulted in less variation between samples (Figure 4). However, this was not evident when categorised according to microscopic OA grading. For native SF biobank samples categorised according to macroscopic grading, the abundance of three uncharacterised proteins and immunoglobulin kappa constant decreased with increasing OA severity. An uncharacterised protein (a member of the superfamily containing a leucine-rich repeat) and ApoA1 abundances both increased initially with OA pathology but then returned towards baseline with greater OA severity. Microscopic OA categorisation for native SF biobank as well as macroscopic and microscopic OA categorisation of native SF HKJC samples did not identify any differentially abundant proteins. The proteomes of native SF HKJC samples, categorised according to synovitis grade, were not separated via PCA (Figure 5). However, the abundance of afamin was identified as differentially abundant between low and high grade synovitis, decreasing with synovitis severity. ProteoMiner™ processing of both biobank and HKJC SF did not identify any differentially abundant proteins or distinct proteome clusters (PCA) for any OA or synovitis categorisations (Figures S7 and S8). However, LC-MS/MS analysis of ProteoMiner™ processed biobank SF identified trends in gelsolin and LBP abundance, decreasing with increasing OA severity (Figure 6). ELISA analysis of gelsolin using the same SF samples that were analysed by LC-MS/MS corroborated this finding and was additionally supported by an independent clinical cohort, with gelsolin abundance lower in clinical OA cases compared to controls. LBP ELISA analysis of both dependent and independent SF samples also supported the trend identified via LC-MS/MS, although statistical significance was not reached. For biobank native and ProteoMiner™ processed SF we identified a trend that ApoA1 increased initially with OA severity and subsequently decreased. This trend was supported by ApoA1 ELISA analysis using dependent SF samples; however statistical significance was not reached.

**Figure 4.**
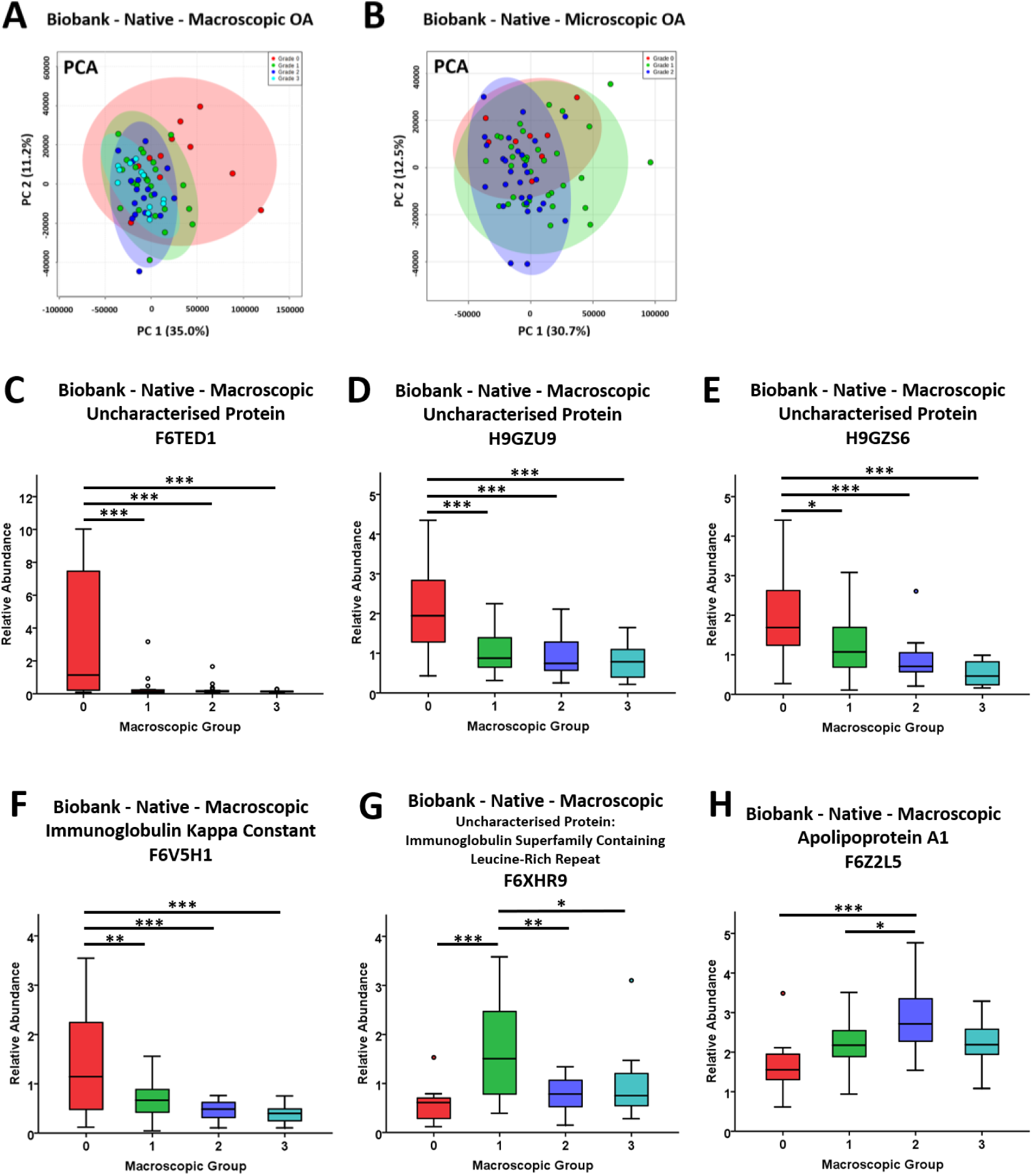
Principal component analysis (PCA) of the biobank native synovial fluid proteome categorised by (A) macroscopic osteoarthritis (OA) grade and (B) microscopic OA grade using LC-MS/MS. (C-H) Differentially expressed proteins when categorised according to macroscopic OA grade. ANOVA: * = p < 0.05, ** = p < 0.01, *** = p < 0.001. Macroscopic OA, n=66, Microscopic OA, n=70.

**Figure 5.**
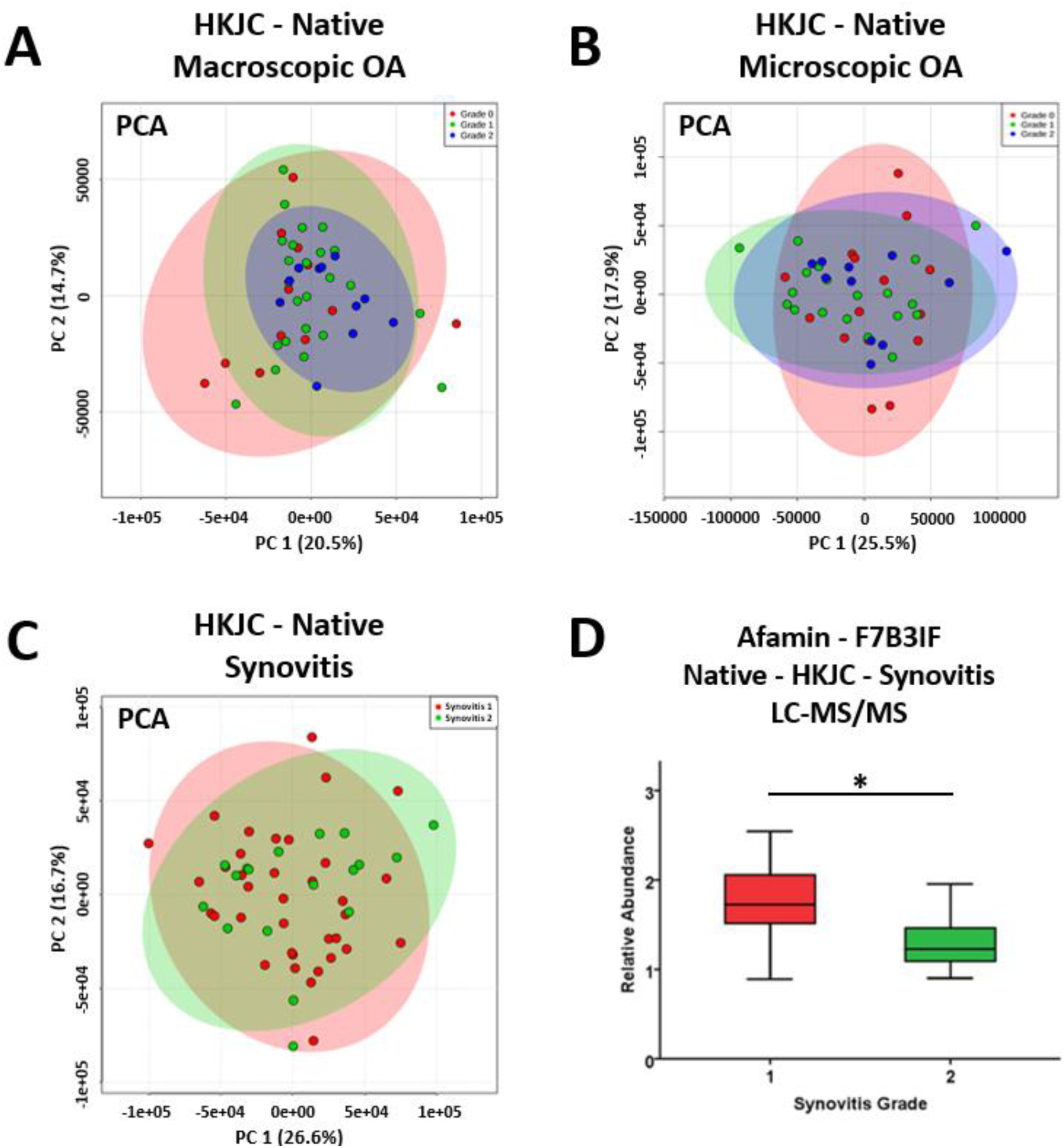
Principal component analysis (PCA) of the Hong Kong Jockey Club (HKJC) native synovial fluid proteome profile categorised by (A) macroscopic osteoarthritis (OA) grade (n=49), (B) microscopic OA grade (n=49) and (C) synovitis grade (n=53) using LC-MS/MS. (D) Differential expression of afamin identified between synovitis grade 1 and 2. t-test: * = p < 0.05.

**Figure 6.**
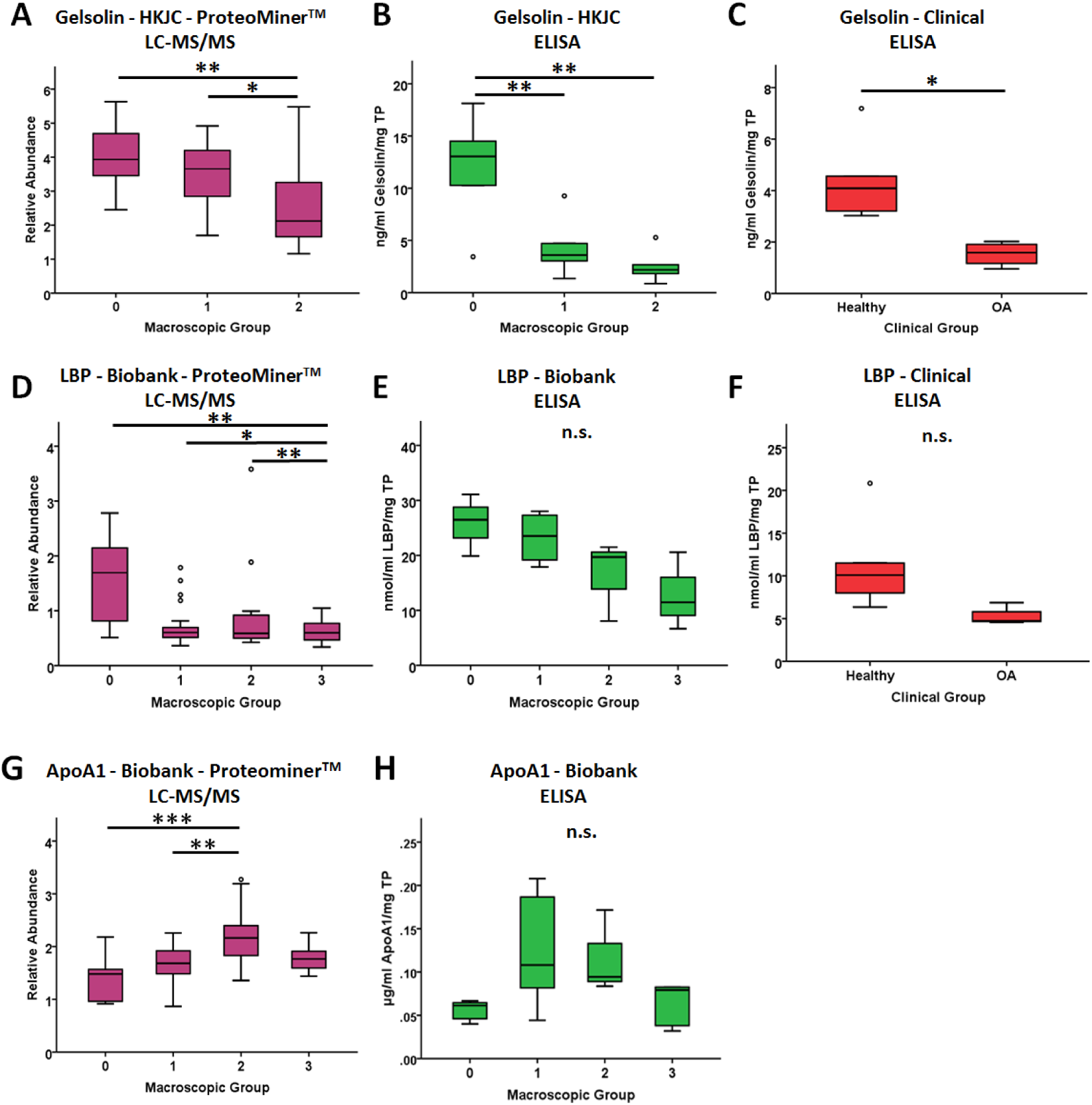
Abundances of gelsolin, lipopolysaccharide binding protein (LBP) and apolipoprotein A1 (ApoA1) within equine synovial fluid via LC-MS/MS (A, D & G) and ELISA validation using dependent (B, E & H) and independent (C & F) synovial fluid samples according to osteoarthritis (OA) grade. LC-MS/MS; HKJC, n=47; Biobank, n=60; ELISA, n=3-6/group. ANOVA and t-tests: * = p < 0.05, ** = p < 0.01, *** = p < 0.001. LC-MS/MS p values corrected for multiple testing within sample, but not for entire dataset.

#### Semi-Tryptic Peptide Profiles

No semi-tryptic peptides arising from ECM proteins were identified which were more highly abundant at more severe OA levels and thus no potential neopeptide biomarkers were identified. Neither for the biobank or HKJC groups were distinct profiles identified between severities of OA (Figure S9). For the HKJC group there was less intra-group variation identified with increased levels of OA and synovitis pathology. However, conversely for the biobank group, samples with no OA pathology showed the least variation.

#### NMR metabolomics and LC-MS/MS Proteomics Integration

For the HKJC dataset, linear regression using Lasso penalisation modelling was unable to select a suitable number of parameters without overfitting the model. However, correlation analysis of all variables (proteins and metabolites) identified 58 significant variables, with a range in correlation from -0.48 to 0.42 (Table S2). Using these selected variables, PCA identified less variation between samples with more severe macroscopic OA scores, although their grouping could not clearly be distinguished from lower scoring samples (Figure 7).

**Figure 7.**
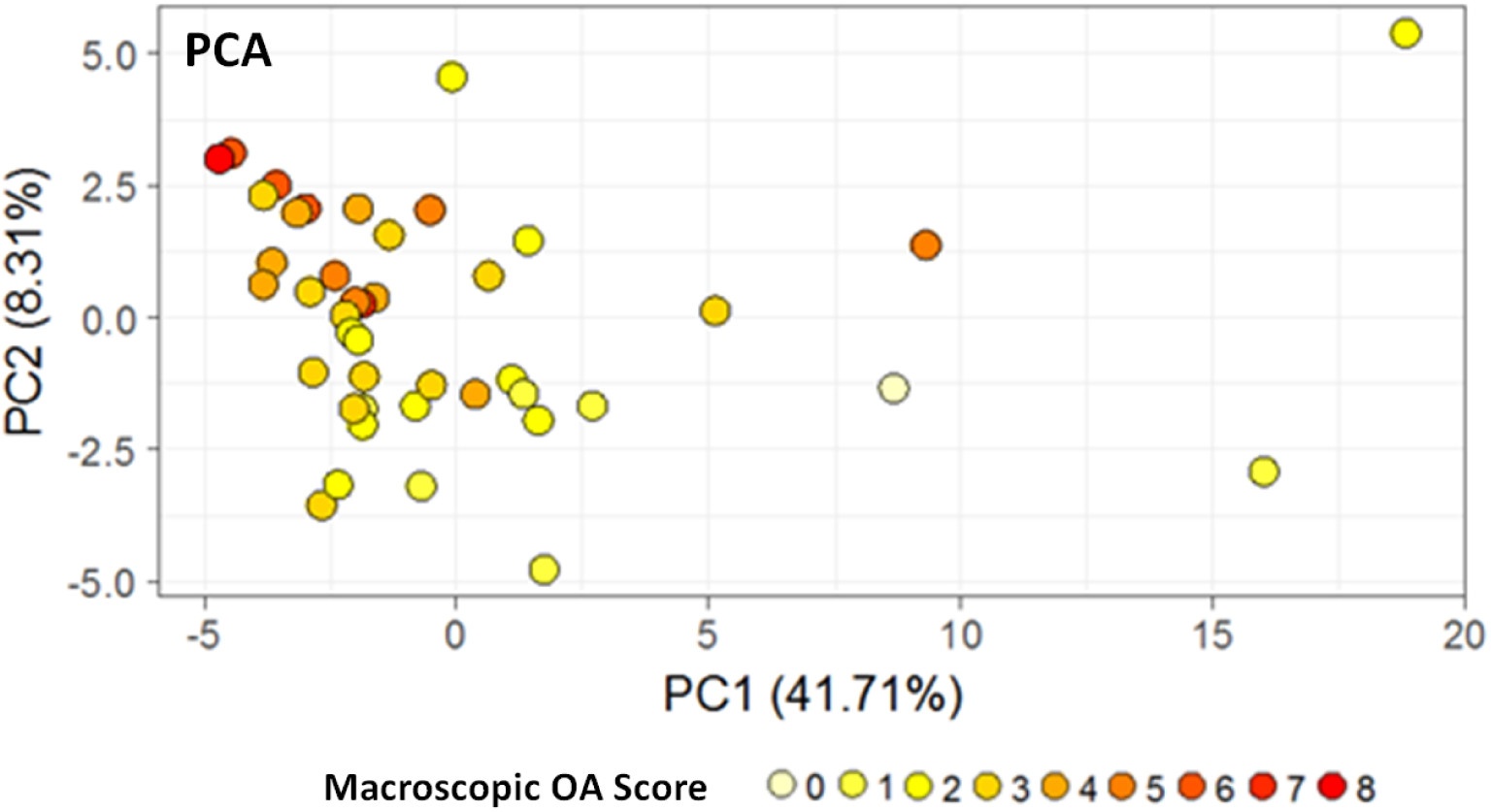
Principal component analysis (PCA) using 58 selected variables (proteins and metabolites) for the Hong Kong Jockey Club synovial fluid combined datasets (n=43) following batch correction, grouped according to macroscopic osteoarthritis score. OA = osteoarthritis.

For the biobank dataset, Lasso penalisation produced a model incorporating 29 variables (Table S3). This produced a good model (R^2^ = 0.82) with PC2 able to discriminate more advanced stages of OA from the earlier stages (Figure 8). This model was predominantly driven by two uncharacterised proteins (H9GZQ9 and F6ZR63) and alanine. Correlation analysis of all variables (proteins and metabolites) identified 32 significant variables, with a range in correlation from -0.44 to 0.49 (Table S4).

**Figure 8.**
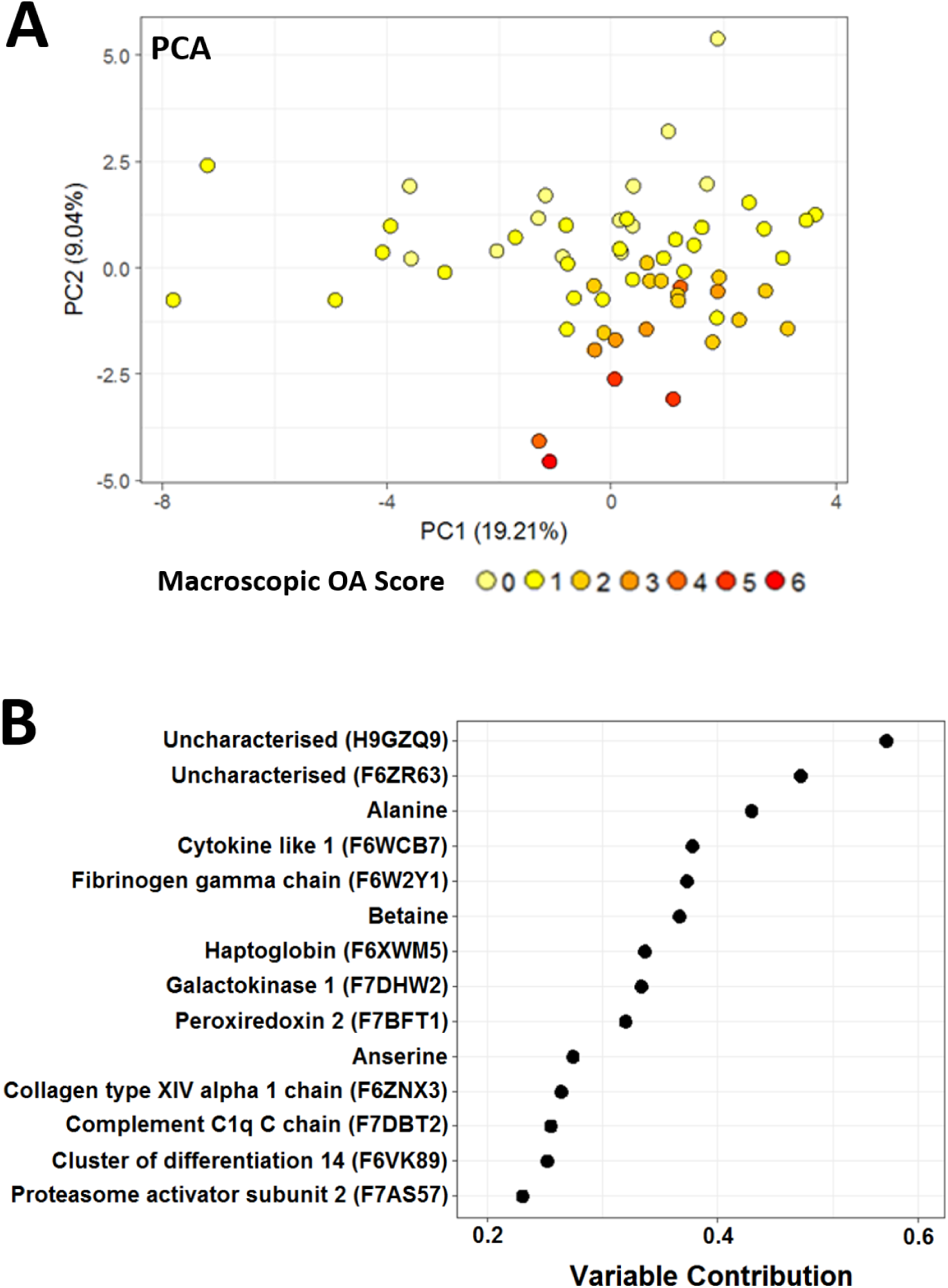
(A) Principal component analysis (PCA) following Lasso model selection using the top 29 variables of influence for biobank synovial fluid, integrating metabolite and protein abundances and grouped according to macroscopic osteoarthritis scoring. (B) Top 14 variables of influence contributing to the model. n=60.

Combining both the biobank and HKJC datasets, including significant protein and metabolite variables, DAPC analysis produced a model whereby OA severity prediction was good for mild OA (90%) although this was reduced significantly for healthy (57%) and severe OA (35%) classifications (Figure 9). Linear discriminant 1 was driven predominantly by neural EGFL like 2, serum albumin and alanine whilst linear discriminant 2 was driven by periostin, gelsolin and myocilin.

**Figure 9.**
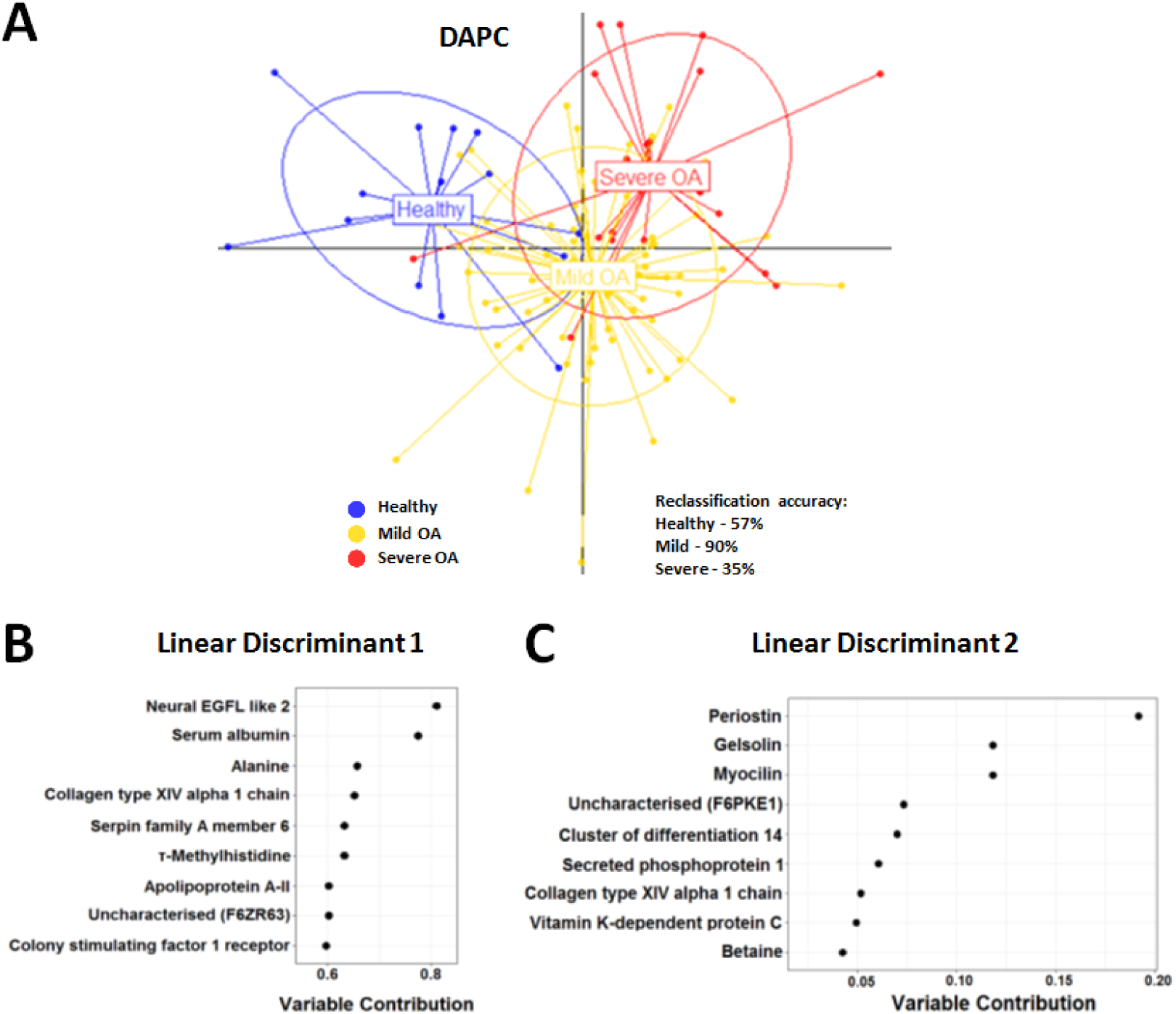
(A) Discriminant analysis of principal components (DAPC) of selected variables (proteins and metabolites) for a combined biobank and Hong Kong Jockey Club synovial fluid dataset (n=103), built using 20 principal components and two linear discriminants. (B) Top 25% of variables for the DAPC model contributing to linear discriminant 1 and (C) linear discriminant 2. Macroscopic osteoarthritis (OA) scoring; Healthy = 0, Mild OA = 1-3, Severe OA = 4-8.

#### Uncharacterised Proteins

BLAST analysis of amino acid sequences of 12 uncharacterised proteins included within this study identified various related characterised proteins. The characterised protein with the highest percentage amino acid sequence similarity for each uncharacterised protein is shown in Table S5.

#### Pathway Analysis

Pathway analysis conducted on proteins and metabolites which were considered significant variables during DAPC modelling, when carrying out NMR metabolomic and LC-MS/MS proteomic dataset integration, identified the complement and coagulation cascades pathway and ABC transporters pathway to be the most represented (Tables 3 and 4). These pathways consisted of 7/96 proteins and 4/9 metabolites included within the pathway analysis respectively.

**Table 3.** Pathway analysis conducted on proteins which were considered significant variables during DAPC modelling, when carrying out NMR metabolomic and LC-MS/MS proteomic dataset integration.

| Pathway ID | Pathway Description | Number of Proteins in Pathway | False Discovery Rate |
| --- | --- | --- | --- |
| 4610 | Complement and coagulation cascades | 7 | 0.00000573 |
| 4145 | Phagosome | 6 | 0.00498 |
| 4390 | Hippo signalling pathway | 5 | 0.0188 |

**Table 4.** Pathway analysis conducted on metabolites which were considered significant variables during DAPC modelling, when carrying out NMR metabolomic and LC-MS/MS proteomic dataset integration.

| Pathway ID | Pathway Description | Number of Metabolites in Pathway |
| --- | --- | --- |
| 2010 | ABC transporters | 4 |
| 4978 | Mineral absorption | 3 |
| 10 | Glycolysis / Gluconeogenesis | 2 |
| 4974 | Protein digestion and absorption | 2 |
| 340 | Histidine metabolism | 2 |
| 4066 | HIF-1 signalling pathway | 2 |
| 4976 | Bile secretion | 2 |
| 4922 | Glucagon signalling pathway | 2 |
| 970 | Aminoacyl-tRNA biosynthesis | 2 |
| 1230 | Biosynthesis of amino acids | 2 |

## Discussion

Equine OA is currently predominantly diagnosed through radiography and clinical examination, however, due to the slow onset of the condition this often leads to substantial pathology of the joint and articular cartilage degradation prior to diagnosis (9). Presently, no equine OA-specific biomarkers have been identified which are able to accurately diagnose and stratify OA with none used to aid an earlier clinical diagnosis (10). Within this study we have investigated equine SF of varying severities of associated OA pathology using both NMR-led metabolomics and MS-based proteomics approaches.

Although both the biobank and HKJC datasets produced a degree of OA stratification, markers of interest were generally distinct. This may be reflective of the more severe OA phenotype demonstrated by the HKJC horses, whilst markers identified within the biobank represent a less severe form of OA. Alternatively, the differences in markers identified may be due to the varying OA aetiologies, with the biobank representing a naturally occurring OA whilst the HKJC horses demonstrate a POD model for subchondral bone mediated OA, which is associated with trauma and overload (23). Additionally, the biobank cohort is on average an older donor group than the HKJC cohort which may also account for differences between the datasets. ^1^H NMR SF spectra were more variable between biobank donors compared to the HKJC donors. This is likely because the biobank is a more heterogeneous population, including age, breed, work loads and diet. The HKJC group is a much more controlled sample set, with all horses housed together, all Thoroughbred racehorses, generally fed and trained similarly and all of a similar age at euthanasia. These differentials may also reflect the different markers of interest identified between the biobank and HKJC groups.

Proteomic analysis of biobank samples identified several proteins which were able to discriminate a healthy phenotype from early OA changes. Although these proteins were largely uncharacterised, BLAST analysis of the amino acid sequences of three uncharacterised proteins, decreasing in abundance with OA severity, identified high levels of similarity to immunoglobulin gamma 1 heavy chain constant region and immunoglobulin kappa chain V-III region MOPC 63 proteins. However, abundances of these uncharacterised proteins could not be validated using antibody methodologies. Recently, a study investigating the SF proteome of a surgery-induced OA model in rabbits identified reduced levels of immunoglobulin heavy chain protein compared to sham controls (69). Immunoglobulins have previously been identified within superficial articular cartilage layers of a proportion of OA patients as well as elevated levels found within the synovial membrane of dogs diagnosed with cranial cruciate ligament rupture (70, 71). Thus, reduction of these immunoglobulins within the SF may be reflective of their translocation to surrounding articular tissues.

Glutamate is an excitatory amino acid neurotransmitter within the central nervous system, although evidence also suggests glutamate operates through intercellular signally cascades, as an autocrine/paracrine factor, in non-neuronal tissues (72). A self-sufficient glutamate signalling machinery has been identified within chondrocytes with a peripheral NMDA receptor proposed to have a role within inflammation and cartilage degradation (73). Within our study, the biobank group showed a reduction in synovial glutamate levels at higher OA severities. Previously, elevated levels of glutamate have been identified within OA SF of humans and within an OA rat models (74–76). However, no increase in glutamate was identified with equine OA SF using ^1^H NMR, although it is not clear whether glutamate was identified during this study (32). The disparity between the lower glutamate levels identified within the biobank group and elevated levels within the literature may be reflective of the subtler OA phenotype exhibited by the biobank samples, and they may be indicative of glutamate levels at an earlier OA severity. However, the HKJC dataset, reflective of a higher grade OA, did not identify differential abundance. Therefore, at this stage the relationship between synovial glutamate abundance and equine OA remains inconclusive.

Creatine is a nonessential amino acid involved in cellular energy metabolism, maintaining cellular adenosine triphosphate (ATP) levels, in particular within the muscle and brain (77). Creatine SF levels increased with OA severity, which is supported by previous human and equine studies which also identified elevated SF creatine in OA SF (32, 78). Approximately 95% of stored creatine is located within skeletal muscle (79). Thus, given the association of muscle atrophy with OA, this elevation in synovial creatine may be reflective of an associated muscle mass loss (80).

Within this study, alanine levels were identified to be increasing in SF in accordance with OA severity, which was also observed by Lacitignolia *et al*. (32). Previously, depleted alanine abundance has been identified within human OA cartilage using high resolution magic angle spinning NMR spectroscopy (81). As alanine is one of the main amino acid residues which constitutes collagen, it may be that the reduction in alanine abundance identified with OA cartilage is resultant of degradation of the cartilage collagen framework, which are subsequently released into the SF resulting in the elevated synovial abundance within this study (81, 82).

Gelsolin (82-84 kDa) is a multifunctional, calcium ion-regulated actin filament severing, capping, and nucleating protein which is involved in the determination of cell shape, secretion and chemotaxis (83, 84). Previous studies of gelsolin on different biological systems have identified gelsolin as a potential predictor of both inflammation and tissue injury (85–87). Within rheumatoid arthritis patients, reduced circulating levels of plasma gelsolin have been identified (88). The authors propose this may be due to gelsolin redistribution to the affected joint space, binding to a plasma factor, reduced production or increased degradation. Within this study, SF gelsolin abundance was identified as decreasing with OA severity. Gelsolin was also identified as having a significant correlation with OA severity and was found to be a highly influential factor when a model combining both biobank and HKJC datasets was developed. These results are supported by a mouse model whereby gelsolin knockout mice resulted in arthritis exacerbation (89). Additionally, within a mouse model of pain and acute inflammation, exogenous delivery of gelsolin was identified to have effective analgesic and anti-inflammatory properties (90). Exogenous gelsolin administration has also been shown to have chondroprotective properties, nullifying the effect of interleukin-1β and OA SF on anabolic gene expression and increased glycosaminoglycan deposition in chondrocytes and protection of the integrity of murine cartilage following intra-articular injection (84). Therefore, gelsolin has potential as a biomarker of equine OA as well as an OA therapeutic target, and potential analgesic.

LBP is an endogenous protein which binds to lipopolysaccharides and catalytically delivers monomeric liposaccharides to cluster of differentiation 14 (CD14) protein (91, 92). Previously, serum and synovial levels of LBP were not identified to be differentially expressed between human degenerative arthropathy patients and control samples (93). However, increased plasma LBP levels have recently been described to predict knee OA progression (94). Within this current study, the trends identified suggest a decreasing SF abundance of LBP with increasing OA severity. It has previously been identified that mononuclear cell activation, induced by lipopolysaccharides, is enhanced by low LBP concentrations (95). Activation of monocytes and macrophages by lipopolysaccharides leads to the secretion of tumour necrosis factor alpha and interleukin-1 beta, two pro-inflammatory cytokines which are central to OA pathogenesis (96, 97). Thus, decreasing synovial LBP levels may have a role in OA development. With CD14 also identified as a key variable within the biobank and HKJC combined model, this provides further support that this pathway is involved in equine OA pathogenesis.

Elevations in synovial ApoA1 were identified in mild and moderate OA within the biobank group although abundance decreased for severe OA. Elevations in OA SF have previously been identified in horses and dogs (39, 98). ApoA1 is the primary protein component of high density lipoproteins (HDLs) and involved in HDL binding to ATP-binding cassette (ABC) transporters as well as being a lecithin cholesterol acyl transferase cofactor (99–102). ApoA1 has previously been found to induce the expression of interleukin-6, MMP-1 and MMP-3 in chondrocytes and synoviocytes through toll-like receptor 4 with the same study identifying a dissociation between the relationship of ApoA1 and HDLs in OA SF (103). When combining datasets within this study, apolipoprotein A-II (ApoA2) was also found to be an important variable.

ApoA2 has previously been identified to be involved in the acute phase response which is associated with reactive amyloid A amyloidosis, via lipoprotein conformational changes, an associated complication of rheumatoid arthritis (104). Thus, this study provides further evidence that OA is a metabolic syndrome with disruption of lipid homeostasis due to alterations to apolipoprotein activity (103).

Synovitis has previously been identified as an important aspect of OA pathogenesis (105–107). This study identified reduced synovial abundance of afamin in high grade synovitis compared to low grade. Similarly, reduced afamin levels have been recorded in equine OA SF compared to healthy joints (39). In a previous study however, elevated levels of afamin were identified within human knee OA SF (108). Afamin is a vitamin E binding glycoprotein and a member of the albumin gene family (109). Afamin forms a 1:1 complex with various hydrophobic Wnt proteins, solubilising the proteins and producing a biologically active complex (110). A growing body of evidence has identified that the Wnt/β-catenin signalling cascade is likely to have a central role within OA pathogenesis (111). Thus, a reduction in synovial abundance of afamin may be reflective of a translocation following Wnt solubilisation to surrounding articular tissues, i.e. the synovium. However, it should be noted that a differential abundance of afamin was not identified when categorised according to OA severity.

For the HKJC the semi-tryptic profile was most consistent between samples in the groups with the most severe OA and synovitis pathology. This suggests, as expected, the OA phenotype is driving the semi-tryptic peptide profile, most likely due to an increase in enzymatic activity leading to ECM degradation fragments (112). However, this was not evident within the biobank SF sample set. This may be because the OA pathology identified within this group was far subtler than that identified within the HKJC samples and thus the level of pathology present was not severe enough to drive a global change within the semi-tryptic peptide profile.

Computational integration of separate ‘omics’ datasets can work synergistically, greatly enhancing the information that can be obtained from separate analyses (113). However, this poses the challenge of developing multi ‘omic’ integration techniques. Previously, a study carried out MS proteomic analysis comparing early and late stage human OA SF and combined this with transcriptomics of articular tissues to identify the source of differentially abundant synovial proteins (114). However, no previous studies have combined NMR metabolomics and MS proteomic datasets.

Integrating the metabolomic and proteomic datasets separately for the biobank and HKJC both identified groupings associated with OA severity. Although correlation values of variables to OA severity were generally low for the HKJC samples, stratification was still evident. Modelling was most robust for the biobank dataset, with the application of a Lasso model, with the model’s strength being the separation of more severe OA grades. However, within this sample set there were relatively few samples which were assigned to the higher OA severity grades compared to lower and is therefore a limitation of the model. Combining all results from both the biobank and HKJC datasets produced a model which was found to be highly accurate in correctly assigning samples within the mild OA group based on their integrated metabolomic and proteomic profiles. This is promising as mild OA recognition is the current unmet clinical need, aiding in early diagnosis and allowing for timely OA interventional management. However, the model is dominated by predominantly mild OA graded samples, and thus this will inevitably introduce bias into the model and make a correct reclassification of mild OA more likely.

Within the combined dataset DAPC model, periostin was the principal driver of linear discriminant 2. Periostin is a 90 kDa matricellular protein which has regulatory functions in cell differentiation, cell adhesion and ECM organisation (115, 116). In human medicine, a positive correlation has been identified between both plasma and SF periostin abundance and knee OA severity (117). *In-vivo* results have identified that interleukin-13 may induce the production of periostin during OA, with periostin subsequently stimulating the production of MMPs within synoviocytes (118). In women, serum periostin levels were found to be associated with both prevalence and the risk of development/progression of knee OA (119). Thus for equids, periostin may also provide a potential therapeutic target and prognostic indicator.

Protein pathway analysis identified the ‘complement and coagulation cascades’ pathway to be altered during OA pathology. Additionally, the uncharacterised protein F6ZR63, the second most important variable of influence driving the biobank Lasso model, was found to have a high level of amino acid sequence similarity to complement factor H-like isoform X1. The complement system is an important component of the innate immune response, encompassing various roles including opsonisation initiation, pathogen phagocytosis, the inflammatory response and terminating within cell lysis (120, 121). However, a growing body of evidence has identified a role of complement activation within OA pathogenesis, which is further supported by this study (120). Complement has been identified to have a role in the degradation of cartilage ECM, synovitis and osteophyte formation, with complement split fragment C3a found to upregulate gene expression of tumour necrosis factor-α and Interleukin-1β, pro-inflammatory cytokines central to OA pathogenesis (97, 120). Therefore, targeting the complement cascade may provide a novel therapeutic target for OA treatment.

Within experimental arthritis models, coagulation and fibrinolysis pathways are known to play a role within disease pathogenesis, with these cascades also found to be activated within both the joint and circulation of degenerative and inflammatory arthropathies (122). Within the synovium of patients diagnosed with OA, the coagulation factor fibrinogen is present throughout the tissue with elevations in tissue factor, a coagulation initiator, identified near endothelial cells (123). Previously, fibrinogen has been identified as a potential target for arthritis therapy, with removal of fibrinogen-leukocyte integrin receptor αMβ2 interactions limiting inflammatory processes whilst maintaining normal fibrinogen coagulation function (124). Thus these pathways may also hold potential as a novel target for OA therapy.

Metabolite pathway analysis identified the ‘ABC transporters’ pathway to be altered during OA pathology. ABC transporters utilise ATP hydrolysis to import or export molecules across cell membranes (125). ABC exporters have an important role in the export of cholesterol, fatty acid and lipids from cells, with dysregulation of this pathway underlying numerous diseases. The Wnt/β-catenin signalling cascade, with a proposed central role within OA pathogenesis, has previously been identified as a regulator of ABC transporters (126). Additionally, the ABC transporter MRP5 has been found to be the principal exporter of hyaluronan from its site of synthesis within the cell to the ECM (127). However, as only four key metabolites were identified within this pathway, caution should be applied when analysing these results to avoid over interpretation.

### Further Work

Several proteins of interest identified within this study were uncharacterised. BLAST amino acid sequence analysis of similar proteins provided further information on their potential function, however this could be taken further by conducting 3D modelling of these proteins to help confirm function and potential interactions with other proteins/molecules. Additionally, developing monoclonal antibodies which are specific to these uncharacterised proteins would allow this methodology to provide an orthologous method to confirm protein abundances. Within the cohorts used in this study, there was an over-representation of mild OA diagnoses opposed to more severe phenotypes. Although this is advantageous in terms of identifying early OA markers, in order to fully stratify OA, the addition of a greater number of donors with higher grade OA would provide a more comprehensive synovial profile of the metabolite and protein profiles at differing OA grades and provide more robust models. As this study adopted a cross-sectional approach, further work of interest would be to conduct repeated measures within a longitudinal study to discover whether the identified markers of interest validate this study’s findings. Also, given the differential abundance of ApoA1 and LBP identified within this study, it would be of interest to further interrogate SF, using both NMR and MS based approaches, to investigate the lipid profile and identify changes during OA progression.

## Conclusions

In conclusion, this study has stratified equine OA using both metabolomic and proteomic SF profiles and identified a panel of markers of interest which may be applicable to grading OA severity. This is also the first study to undertake computational integration of NMR metabolomic and LC-MS/MS proteomic datasets of any biological system.

## Acknowledgements

Dr James Anderson was funded through a Horse Trust PhD studentship (G1015) and Dr Mandy Peffers funded through a Wellcome Trust Intermediate Clinical Fellowship (107471/Z/15/Z). Software licenses for data analysis used in the Shared Research Facility for NMR metabolomics were funded by the Medical Research Council (MRC) Clinical Research Capabilities and Technologies Initiative (MR/M009114/1). ELISA protein validation experiments were funded by the Institute of Veterinary Science, University of Liverpool. This work was also supported by the MRC and Versus Arthritis as part of the Medical Research Council Versus Arthritis Centre for Integrated Research into Musculoskeletal Ageing (CIMA) [MR/R502182/1] and a Technology Directorate Voucher (Faculty of Health and Life Sciences, University of Liverpool). The MRC Versus Arthritis Centre for Integrated Research into Musculoskeletal Ageing is a collaboration between the Universities of Liverpool, Sheffield and Newcastle. The authors would like to thank Professor Christopher Riggs, Hong Kong Jockey Club, for providing samples for analysis, staff at both the F Drury and Sons Abattoir, Swindon and Hong Kong Jockey Club for assistance in sample collection and processing, Ms Valerie Tilston for preparation of histology slides, Mr Phillip Dyer for assisting with ProteoMiner™ column processing, members of the Centre for Protein Research, University of Liverpool, including Professor Rob Beynon and Dr Philip Brownridge for guidance and access to their mass spectrometry facilities, Mr Jake Ellis, Cardiff University, for undertaking NMR file depositions and Dr Ben McDermott, Dr Elizabeth Barr, Ms Amy McDermott, Mr Aibek Smagul and Mr Colin Anderson for conducting pathology scoring.

## Ethics

Hong Kong Jockey Club samples were collected under the regulations of the Hong Kong Jockey Club with owner consent. Abattoir samples were collected as a by-product of the agricultural industry, and hence ethical approval was not required. Ethical collection of excess clinical equine synovial fluid from live horses and collection of post mortem equine synovial fluid was approved by The University of Liverpool Veterinary Research Ethics Committee, approval references VREC175 and VREC561.

## Data Availability

1D ^1^H NMR spectra, together with annotated metabolite HMDB IDs and annotation level, is available within the MetaboLights repository (www.ebi.ac.uk/metabolights/MTBLS1645) (128).

Due to the COVID-19 pandemic, we are unfortunately unable to upload mass spectrometry proteomics data at present.

## Abbreviations

ADAMTS: A disintegrin and metalloproteinase with thrombospondin motifs
ApoA1: Apolipoprotein A1
ApoA2: Apolipoprotein A-II
BLAST: Basic local alignment search tool
CPMG: Carr-Purcell-Meiboom-Gill
COMP: Cartilage oligomeric matrix protein
CD14: Cluster of differentiation 14
DAPC: Discriminant analysis of principal components
ECM: Extracellular matrix
FDR: False discovery rate
H & E: Haematoxylin and eosin
HDL: High density lipoprotein
HKJC: Hong Kong Jockey Club
Lasso: Least absolute shrinkage and selection operator
LBP: Lipopolysaccharide binding protein
MMP: Matrix metalloproteinase
MSI: Metabolomics Standards Initiative
MCP: Metacarpophalangeal
MTP: Metatarsophalangeal
OA: Osteoarthritis
POD: Palmar/plantar osteochondral disease
PFA: Paraformaldehyde
PCA: Principal component analysis
PQN: Probabilistic quotient normalisation
Saf O: Safranin O
SF: Synovial fluid
TIC: Total ion current.

