## Supplemental Data for "Metabolomic and Proteomic Stratification of Equine Osteoarthritis"

**Contents**

**Liquid Chromatography Tandem Mass Spectrometry - Detailed Methods**

**Figure S1.** Protocol for synovial fluid (SF) collection and processing prior to NMR metabolomic and LC-MS/MS proteomic analysis.

**Figure S2.** Subchondral bone/cartilage wedge taken at post mortem from the distal articular condylar surface of metacarpal III, lateral to the sagittal ridge.

**Figure S3.** Representative ion chromatograms for native synovial fluid following 16hr + 2hr trypsin digestion using a 90 min liquid chromatography (LC) gradient,

ProteoMiner$^{TM}$ processed synovial fluid following a 4hr Lys-C + 4hr trypsin digestion using a 60 min LC gradient and ProteoMiner$^{TM}$ processed synovial fluid following a 4hr Lys-C + 16hr + 2hr trypsin digestion using a 120 min LC gradient.

**Figure S4.** Principal component analyses of biobank and Hong Kong Jockey Club equine synovial fluid NMR metabolomes and LC-MS/MS proteomes before and after application of a COMBAT batch correction.

**Figure S5.** Correlation between microscopic and macroscopic osteoarthritis grading scores for biobank and Hong Kong Jockey Club cohorts.

**Figure S6.** Quantile plots of biobank and Hong Kong Jockey Club synovial fluid NMR spectra.

**Figure S7.** Principal component analysis of biobank ProteoMiner$^{TM}$ processed (16hr + 2hr trypsin digestion) synovial fluid proteome categorised by macroscopic OA grade and microscopic OA grade using LC-MS/MS.

**Figure S8.** Principal component analysis of the Hong Kong Jockey Club ProteoMiner$^{TM}$ processed (16hr + 2hr trypsin digestion) synovial fluid proteome categorised by macroscopic OA grade, microscopic OA grade and synovitis grade using LC-MS/MS.

**Figure S9.** Principal component analysis (PCA) of equine synovial fluid semi-tryptic peptide profiles grouped according to microscopic osteoarthritis and macroscopic osteoarthritis severity for the biobank cohort and microscopic osteoarthritis, macroscopic osteoarthritis and synovitis severity for the Hong Kong Jockey Club.

**Table S1.** Age comparison between the biobank and Hong Kong Jockey Club (HKJC) cohorts.

**Table S2.** Correlation of each variable (proteins and metabolites) to macroscopic OA score for the Hong Kong Jockey Club synovial fluid integrated dataset.

**Table S3.** Selected variables (proteins and metabolites) used to produce Lasso model for biobank synovial fluid.

**Table S4.** Correlation of each variable (proteins and metabolites) to macroscopic OA score for the biobank synovial fluid integrated dataset.

**Table S5.** BLAST analysis of amino acid sequences of uncharacterised proteins included within this study, identifying the characterised protein with the highest percentage amino acid sequence similarity for each uncharacterised protein.

**Table S6.** Macroscopic osteoarthritis scoring of distal metacarpal III for the University of Liverpool equine biobank.

**Table S7.** Microscopic osteoarthritis scoring of distal metacarpal III for the University of Liverpool equine biobank.

**Table S8.** Macroscopic osteoarthritis scoring of distal metacarpal III or metatarsal III for the Hong Kong Jockey Club sample set.

**Table S9.** Microscopic osteoarthritis scoring of distal metacarpal III or metatarsal III for the Hong Kong Jockey Club sample set.

**Table S10.** Synovitis scoring of distal metacarpal III or metatarsal III for the Hong Kong Jockey Club sample set.

**Table S11.** Synovial fluid metabolite and protein abundances following batch corrections for the University of Liverpool equine biobank (separate excel file)

**Table S12.** Synovial fluid metabolite and protein abundances following batch corrections for the Hong Kong Jockey Club (separate excel file)

**Table S13.** Semi-tryptic peptide abundances for the University of Liverpool equine biobank and Hong Kong Jockey Club sample sets (separate excel file)

**Table S14.** Mass spectrometry data of peptides and proteins identified for all native synovial fluid samples (separate excel file)

**Table S15.** Mass spectrometry data of peptides and proteins identified for all ProteoMiner$^{TM}$ 4 hr Lys-C + 16 hr + 2 hr trypsin digested synovial fluid samples (separate excel file)

**Liquid Chromatography Tandem Mass Spectrometry - Detailed Methods**

Tryptic digests were diluted 5-fold in 0.1% (v/v) trifluoroacetic acid (TFA) and 3% (v/v) acetonitrile and analysed individually, in a random order, via liquid chromatography tandem mass spectrometry (LC-MS/MS) using a 60, 90 or 120 min liquid chromatography (LC) gradient as stated. A Q Exactive$^{TM}$ HF quadrupole-Orbitrap mass spectrometer (Thermo Scientific, Hemel Hempstead, UK) coupled to a Dionex Ultimate 3000 RSLC nano-liquid chromatograph (Thermo Scientific) was used for data-dependent LC-MS/MS analyses. Digests were loaded onto a trapping column (Acclaim PepMap 100, C18, 20 mm x 75 µm) using a loading buffer of 0.1% (v/v) TFA and 2% (v/v) acetonitrile in water for 3 min at a flow rate of 5 µl min$^{-1}$. The trapping column was then set in-line with an analytical column (Easy-Spray PepMap® C18, 15 cm x 75 µm, 2 µm) with peptide elution carried out using a linear gradient of 96.2% A (0.1% (v/v) formic acid):3.8% B (0.1 % (v/v) formic acid in water:acetonitrile (80:20) (v/v)) to 50% A:50% B over 30, 60 0r 90 min at a flow rate of 300 nl min$^{-1}$, followed by washing at 1 % A:99% B for 5 min and re-equilibration of the column to starting conditions. The Q Exactive$^{TM}$ was operated in data dependent positive (ESI+) mode with survey scans between *m/z* 300-2000 acquired at a mass resolution of 70,000 (full width at half maximum) at *m/z* 200 after accumulation of ions to $1x10^6$ target value based on predictive automatic gain control values from the previous full scan. The 10 most intense precursor ions with charge states of between 2+ and 5+ were selected for MS/MS with an isolation window of 2 *m/z* units. Higher-energy collisional dissociation (HCD) was used to fragment peptides using normalised collision energy of 30% with a maximum injection time of 100 ms. Dynamic exclusion of *m/z* values to prevent repeated fragmentation of the same peptide was used with an exclusion time of 20 s.

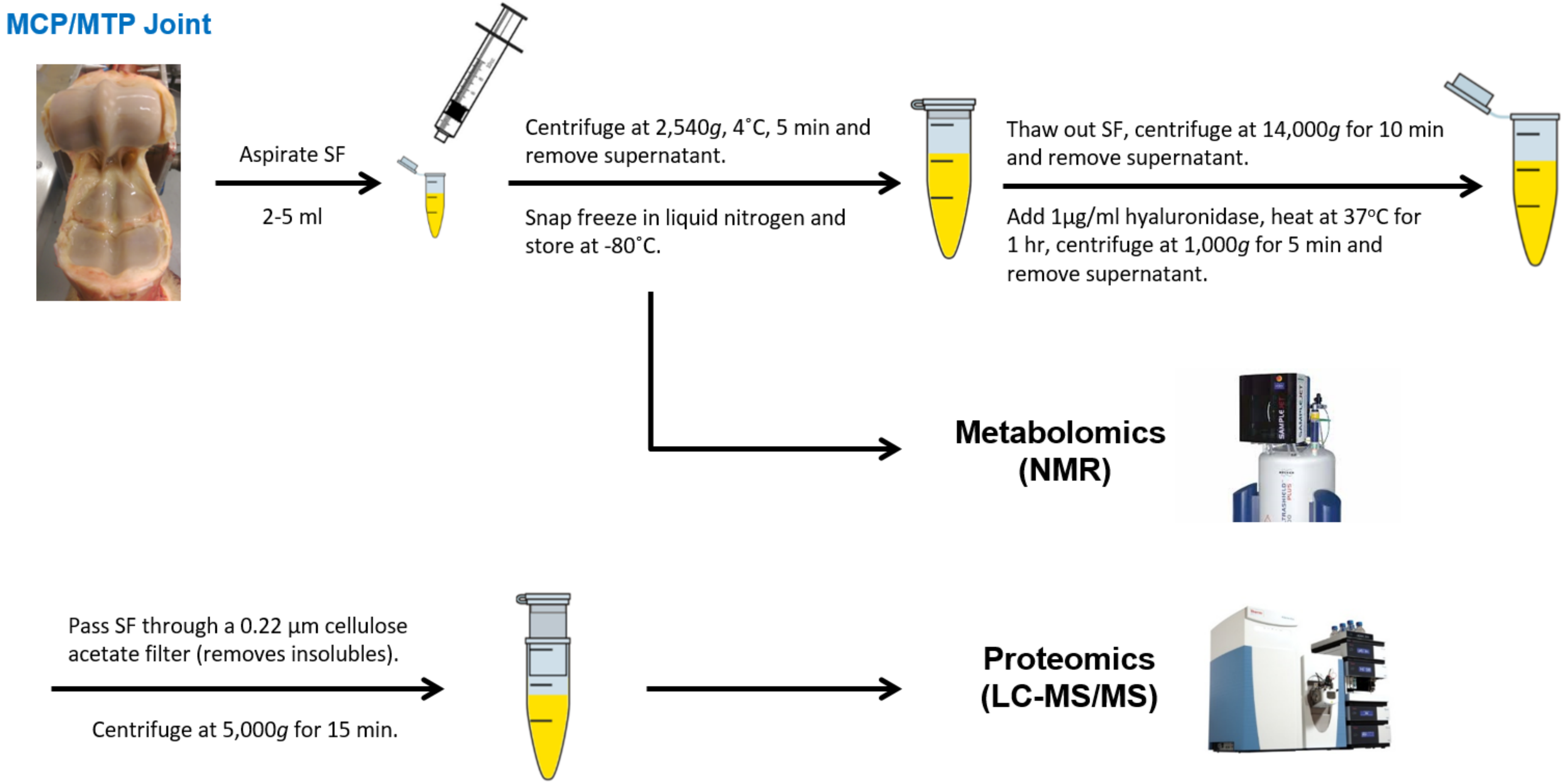

**Figure S1.** Protocol for synovial fluid (SF) collection and processing prior to NMR metabolomic and LC-MS/MS proteomic analysis. MCP = metacarpophalangeal, MTP = metatarsophalangeal.

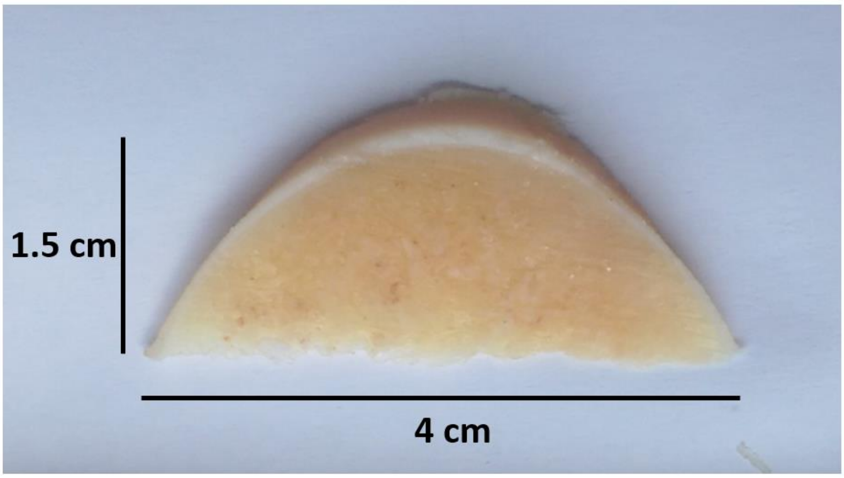

**Figure S2.** Subchondral bone/cartilage wedge taken at post mortem from the distal articular condylar surface of metacarpal III, lateral to the sagittal ridge.

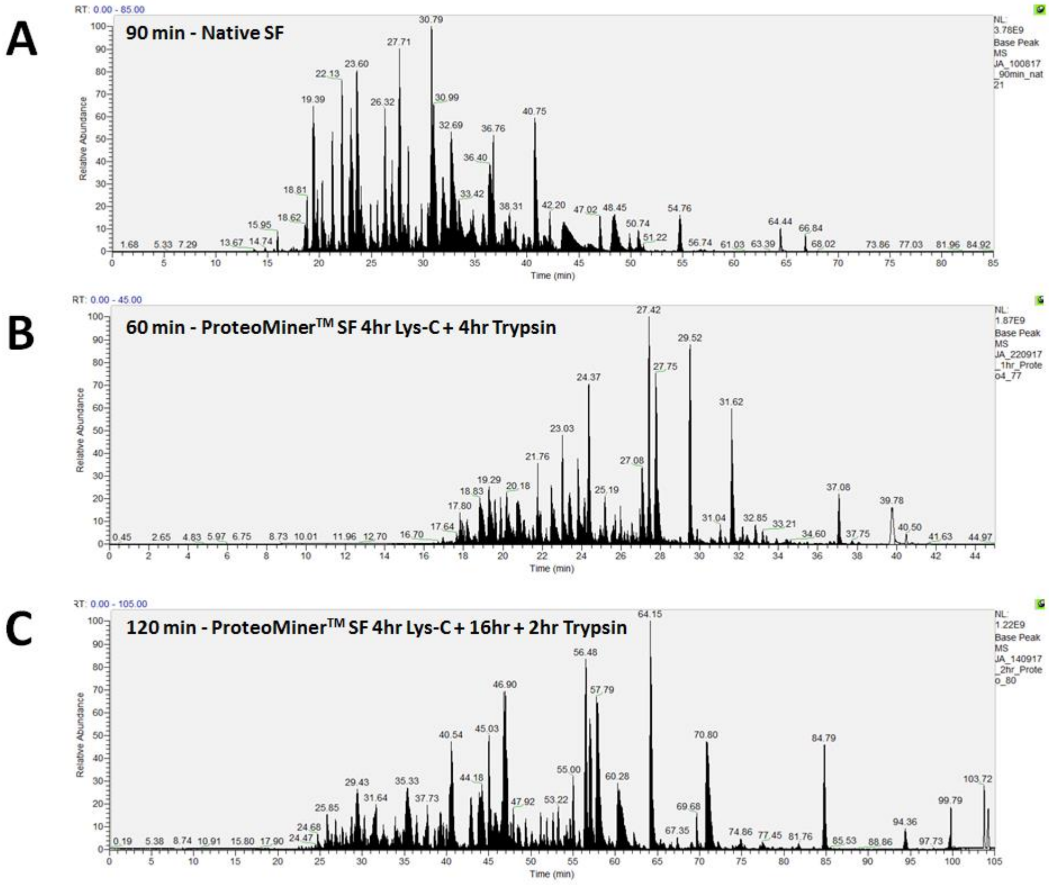

**Figure S3.** Representative ion chromatograms for (A) native synovial fluid (SF) following 16hr + 2hr trypsin digestion using a 90 min liquid chromatography (LC) gradient, (B) ProteoMiner$^{TM}$ processed SF following a 4hr Lys-C + 4hr trypsin digestion using a 60 min LC gradient and (C) ProteoMiner$^{TM}$ processed SF following a 4hr Lys-C + 16hr + 2hr trypsin digestion using a 120 min LC gradient.

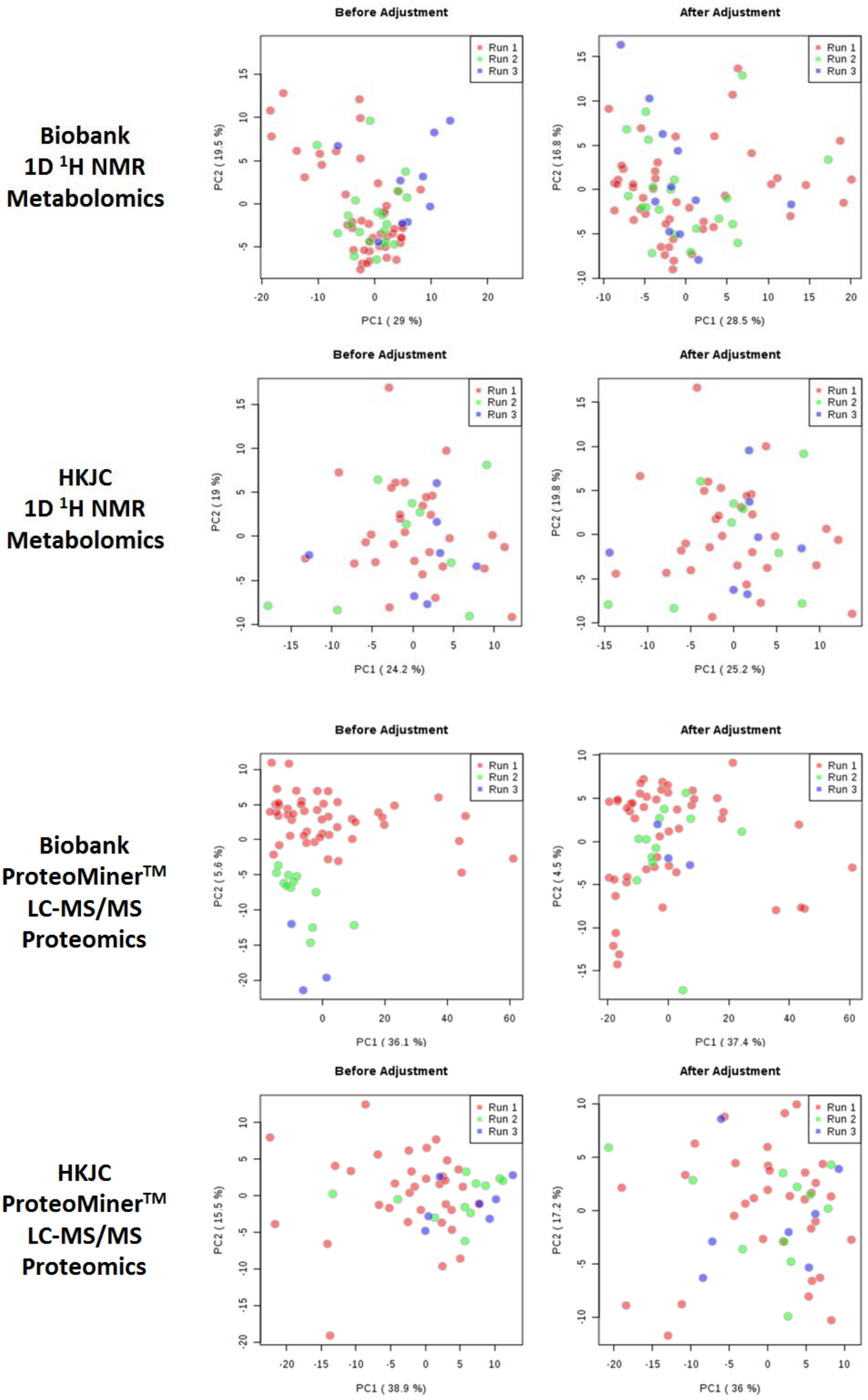

**Figure S4.** Principal component analyses of biobank and Hong Kong Jockey Club (HKJC) equine synovial fluid NMR metabolomes and LC-MS/MS proteomes before and after application of a COMBAT batch correction. biobank; metabolomics, n=76, proteomics, n=70, HKJC; metabolomics, n=56, proteomics, n=53.

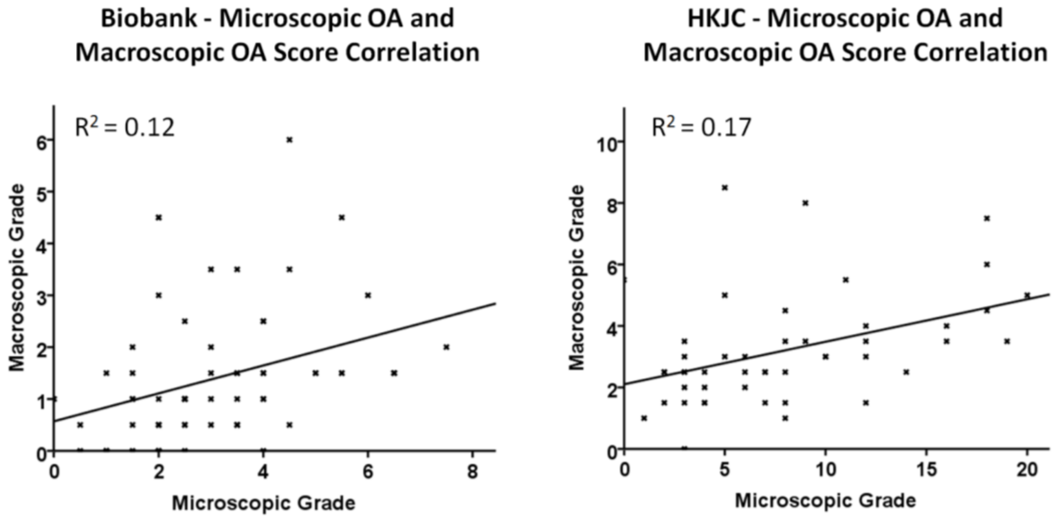

**Figure S5.** Correlation between microscopic and macroscopic osteoarthritis (OA) grading scores for biobank (n=41) and Hong Kong Jockey Club (HKJC, n=41) cohorts.

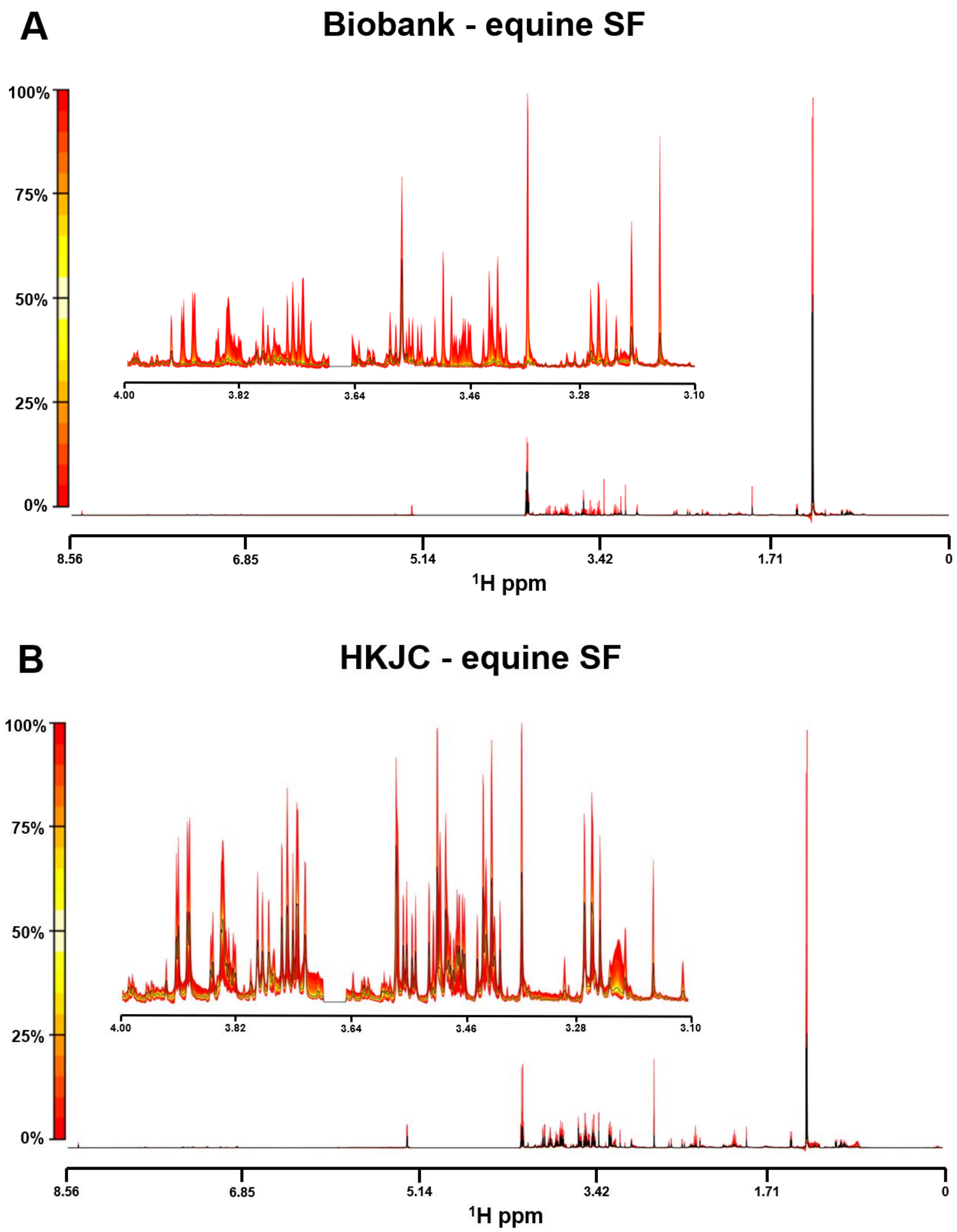

**Figure S6.** Quantile plots of (A) biobank (n=83) and (B) Hong Kong Jockey Club (HKJC, n=52) synovial fluid (SF) NMR spectra. The median spectra are depicted by a black line, with variation from the median spectral plot depicted by a yellow to red scale. The full spectral range is shown (8.56-0 ppm) with a more detailed region inset (4-3.1 ppm). Spectral regions 3.681-3.643 ppm and 1.201-1.162 ppm have been removed due to ethanol contamination.

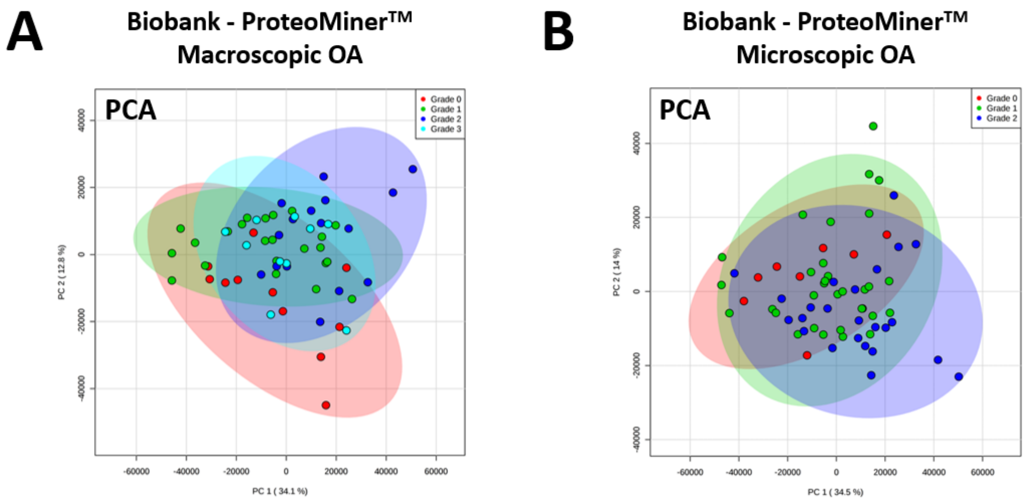

**Figure S7.** Principal component analysis of biobank ProteoMiner™ processed (16hr + 2hr trypsin digestion) synovial fluid proteome categorised by (A) macroscopic OA grade (n=60) and (B) microscopic OA grade (n=64) using LC-MS/MS.

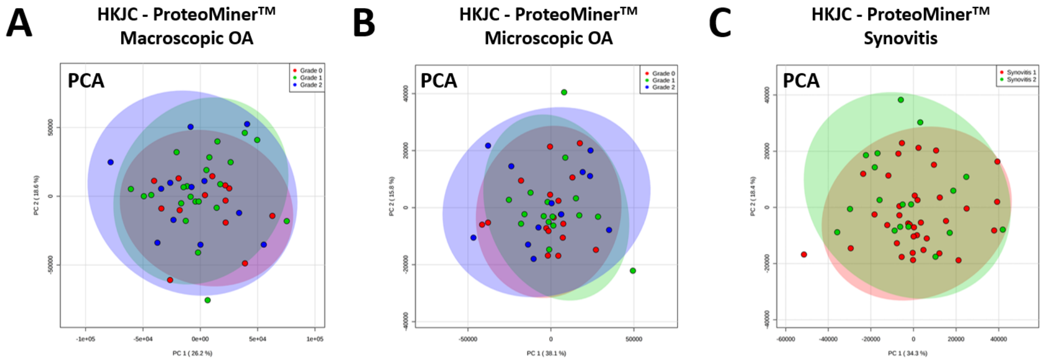

**Figure S8.** Principal component analysis of the Hong Kong Jockey Club (HKJC) ProteoMiner™ processed (16hr + 2hr trypsin digestion) synovial fluid proteome categorised by (A) macroscopic OA grade (n=47), (B) microscopic OA grade (n=45) and (C) synovitis grade (n=53) using LC-MS/MS.

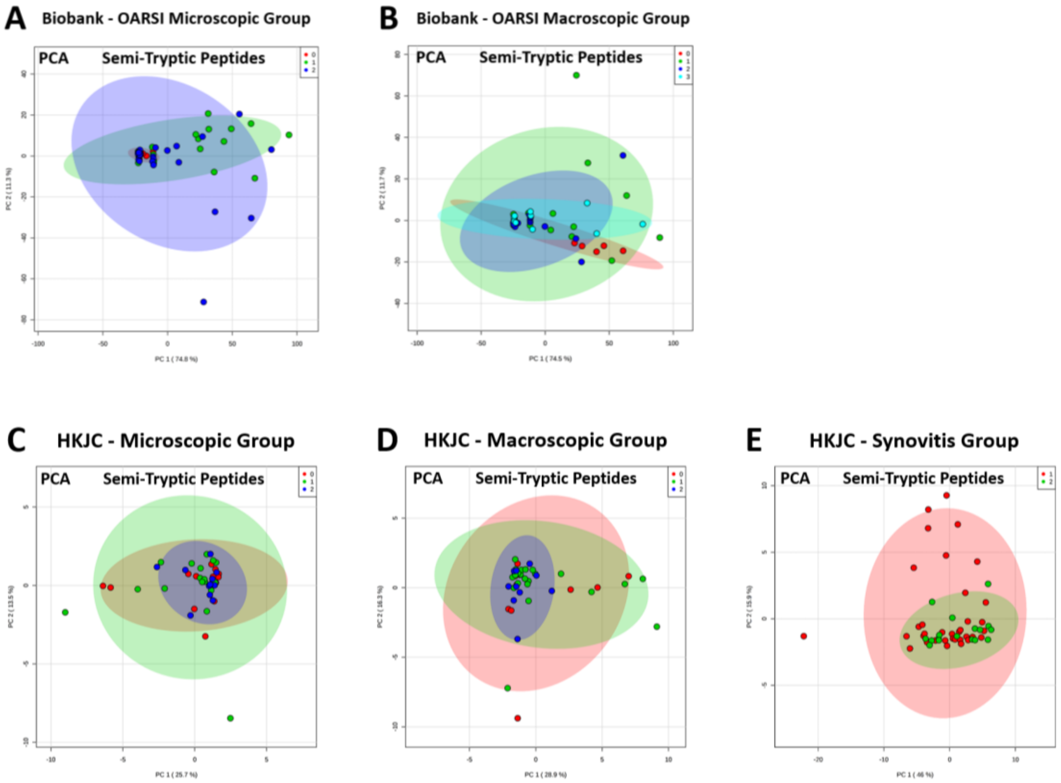

**Figure S9.** Principal component analysis (PCA) of equine synovial fluid semi-tryptic peptide profiles grouped according to (A) microscopic osteoarthritis (n=62) and (B) macroscopic osteoarthritis (n=59) severity for the biobank cohort and (C) microscopic osteoarthritis (n=46), (D) macroscopic osteoarthritis (n=46) and (E) synovitis (n=53) severity for the Hong Kong Jockey Club (HKJC).

**Table S1.** Age comparison between the biobank and Hong Kong Jockey Club (HKJC) cohorts.

| | Biobank | HKJC |
|---|---|---|
| **Number of donors** | 83 | 58 |
| **Mean age (years)** | 14.3 | 6.6 |
| **Median age (years)** | 16 | 7 |
| **Standard deviation (years)** | 7.6 | 1.9 |
| **t-test (mean age)** | $p = 6.22\ e^{-12}$ | |

**Table S2.** Correlation of each variable (proteins and metabolites) to macroscopic OA score for the Hong Kong Jockey Club synovial fluid integrated dataset. $p < 0.05$.

| Variable | Correlation | Permutation p value | Characterisation |
|---|---|---|---|
| P60708 | -0.44 | 0 | Actin, cytoplasmic 1 |
| HMDB00001 | 0.37 | 0 | τ-Methylhistidine |
| F6UL68 | -0.45 | 0 | Transthyretin |
| F6VCB4 | -0.36 | 0 | Histone H3 |
| F6VS95 | -0.48 | 0 | Protein disulfide isomerase family A member 6 |
| F6W3T1 | -0.38 | 0 | L-lactate dehydrogenase |
| F6YNM7 | -0.36 | 0 | Prolyl endopeptidase |
| F6YRC5 | -0.39 | 0 | IQ motif containing GTPase activating protein 1 |
| F6YRE0 | -0.36 | 0 | Tubulin alpha chain |
| F6ZMJ5 | -0.40 | 0 | ATPase $H^+$ transporting V1 subunit E2 |
| F7ABC9 | -0.38 | 0 | Fibulin-1 |
| F7D8W6 | -0.43 | 0 | Guanylate binding protein 2 |
| F7DNT0 | -0.42 | 0 | Tubulin alpha chain |
| F7DYV0 | -0.43 | 0 | Galectin |
| A2Q0Z0 | -0.28 | 0.01 | Elongation factor 1-alpha 1 |
| Q28372 | -0.35 | 0.01 | Gelsolin |
| F6T0P6 | -0.36 | 0.01 | GC, vitamin D binding protein |
| F6UZI2 | -0.35 | 0.01 | Coagulation factor XIII A chain |
| F6V6L8 | -0.36 | 0.01 | IQ motif containing GTPase activating protein 2 |
| F6YIU8 | -0.39 | 0.01 | Methylenetetrahydrofolate dehydrogenase, cyclohydrolase and formyltetrahydrofolate synthetase 1 |
| F6YXS1 | 0.37 | 0.01 | Receptor protein-tyrosine kinase |
| F6ZFH9 | -0.39 | 0.01 | Tyrosine 3-monooxygenase/tryptophan 5-monooxygenase activation protein gamma |
| F7AYC1 | -0.31 | 0.01 | Secreted phosphoprotein 1 |

| | | | |
|---|---|---|---|
| F7BNQ2 | -0.36 | 0.01 | Complement component 4 binding protein alpha |
| F7BQD6 | 0.35 | 0.01 | Complement C1s |
| F7CYG2 | -0.38 | 0.01 | Tyrosine 3-monooxygenase/tryptophan 5-monooxygenase activation protein eta |
| HMDB00062 | 0.35 | 0.02 | Carnitine |
| HMDB00190 | -0.33 | 0.02 | Lactate |
| F6QXN5 | -0.34 | 0.02 | Transgelin |
| F6S0P5 | -0.34 | 0.02 | ADP ribosylation factor 4 |
| F6S2C3 | -0.33 | 0.02 | Cathepsin Z |
| F6UN85 | -0.40 | 0.02 | Carboxylic ester hydrolase |
| F6Y0G5 | 0.42 | 0.02 | Periostin |
| F6Y4J0 | -0.31 | 0.02 | ATPase $H^+$ transporting V1 subunit B2 |
| F6Z8W0 | -0.36 | 0.02 | WD repeat domain 1 |
| F7APS1 | -0.27 | 0.02 | Uncharacterised |
| F7BM69 | -0.31 | 0.02 | Uncharacterised |
| F7BTK9 | -0.36 | 0.02 | Family with sequence similarity 129 member A |
| F7DA17 | -0.39 | 0.02 | Transglutaminase 2 |
| F7DEW5 | -0.31 | 0.02 | Coronin |
| F7DXM5 | 0.32 | 0.02 | Uncharacterised |
| HMDB00161 | 0.29 | 0.03 | L-Alanine |
| F6PKE1 | -0.33 | 0.03 | Uncharacterised |
| F6QKR7 | -0.38 | 0.03 | Myocilin |
| F6RCA8 | -0.34 | 0.03 | Peroxiredoxin 5 |
| F6YV40 | -0.31 | 0.03 | Glyceraldehyde-3-phosphate dehydrogenase |
| F6YV53 | -0.31 | 0.03 | Actinin alpha 4 |
| HMDB00122 | 0.30 | 0.04 | D-Glucose |
| Q95182 | -0.34 | 0.04 | Major allergen Equ c 1 |
| F6RMN2 | -0.34 | 0.04 | Drebrin like |
| F6SN52 | -0.34 | 0.04 | Eukaryotic translation initiation factor 5A |
| F6SP02 | -0.33 | 0.04 | Tyrosine 3-monooxygenase/tryptophan 5-monooxygenase activation protein theta |
| F6T201 | -0.29 | 0.04 | Chondroitin sulfate proteoglycan 4 |
| F6UF75 | -0.33 | 0.04 | Protein S100 |
| F6WWW7 | -0.32 | 0.04 | Capping actin protein of muscle Z-line alpha subunit 1 |
| F6XEB4 | -0.36 | 0.04 | Tyrosine 3-monooxygenase/tryptophan 5-monooxygenase activation protein zeta |
| F7B320 | -0.32 | 0.04 | Dihydropyrimidinase like 2 |
| H9GZU9 | -0.30 | 0.04 | Uncharacterised |

**Table S3.** Selected variables (proteins and metabolites) used to produce Lasso model for biobank synovial fluid.

| Variable | Description |
|---|---|
| HMDB00194 | Anserine |
| F7A1W7 | Apolipoprotein C2 |
| F6XUJ2 | ATP synthase subunit alpha |
| HMDB00043 | Betaine |
| F6TK05 | Chaperonin containing TCP1 subunit 2 |
| F6VK89 | Cluster of differentiation 14 |
| F6ZNX3 | Collagen type XIV alpha 1 chain |
| F7DBT2 | Complement C1q C chain |
| F7BQD6 | Complement C1s |
| F6WCB7 | Cytokine like 1 |
| F6W2Y1 | Fibrinogen gamma chain |
| F7DHW2 | Galactokinase 1 |
| F7BKK5 | Glutathione S-transferase |
| F6XWM5 | Haptoglobin |
| HMDB00161 | L-Alanine |
| HMDB00883 | L-Valine |
| F6X485 | Myosin heavy chain 9 |
| F7BJV0 | Myosin IC |
| F7BFT1 | Peroxiredoxin 2 |
| F7AS57 | Proteasome activator subunit 2 |
| F6XWJ6 | RUN and FYVE domain containing 1 |
| Q2QLB2 | Testin |
| F7CS04 | Tropomyosin 1 |
| F6S2I3 | Uncharacterised |
| F6TED1 | Uncharacterised |
| F6ZR63 | Uncharacterised |
| H9GZQ9 | Uncharacterised |
| F6RYX0 | Vacuolar protein sorting-associated protein 29 |
| F6X667 | Vitamin K-dependent protein C |

**Table S4.** Correlation of each variable (proteins and metabolites) to macroscopic OA score for the biobank synovial fluid integrated dataset. $p < 0.05$.

| Variable | Correlation | Permutation p value | Characterisation |
|---|---|---|---|
| H9GZS6 | -0.44 | 0 | Uncharacterised |
| F7BFT1 | -0.43 | 0 | Peroxiredoxin 2 |
| F6Q4N3 | -0.41 | 0 | Neural EGFL like 2 |
| F6WCB7 | -0.40 | 0 | Cytokine like 1 |
| F6ZR63 | -0.38 | 0 | Uncharacterised |
| H9GZU9 | -0.37 | 0 | Uncharacterised |
| F6RRV1 | -0.35 | 0 | Fetuin B |
| P35747 | 0.33 | 0 | Serum albumin |
| Q28369 | 0.34 | 0 | Retinol-binding protein 4 |
| F6X667 | 0.34 | 0 | Vitamin K-dependent protein C |
| F7A1W7 | 0.49 | 0 | Apolipoprotein C2 |
| H9GZV0 | -0.34 | 0.01 | Uncharacterised |
| F6SP11 | -0.32 | 0.01 | Uncharacterised |
| F6ZRF6 | -0.31 | 0.01 | Serpin family A member 7 |
| F6QF58 | -0.31 | 0.01 | 60S ribosomal protein L6 |
| H9GZQ9 | -0.29 | 0.01 | Uncharacterised |
| F7ASE1 | 0.32 | 0.01 | Interleukin 1 receptor accessory protein |
| F6RM73 | 0.36 | 0.01 | Apolipoprotein A-II |
| F7DRS2 | -0.33 | 0.02 | Serpin family A member 6 |
| F6WLX9 | -0.29 | 0.02 | tRNA-splicing ligase RtcB homolog |
| F6XPL9 | -0.27 | 0.02 | Obg-like ATPase 1 |
| F6XI92 | -0.27 | 0.02 | Bridging integrator 2 |
| F6YV40 | 0.21 | 0.02 | Glyceraldehyde-3-phosphate dehydrogenase |
| F7DBT2 | 0.27 | 0.02 | Complement C1q C chain |
| F6PRI5 | 0.28 | 0.02 | Carboxylic ester hydrolase |
| F6T0P6 | 0.32 | 0.02 | GC, vitamin D binding protein |
| F6UCZ2 | -0.24 | 0.03 | Translin |
| HMDB00562 | 0.30 | 0.03 | Creatinine |
| F6XWJ6 | -0.32 | 0.04 | RUN and FYVE domain containing 1 |
| F6V1V8 | -0.27 | 0.04 | Cluster of differentiation 109 |
| F7BA94 | -0.20 | 0.04 | 40S ribosomal protein S3 |
| F7AS57 | 0.24 | 0.04 | Proteasome activator subunit 2 |

**Table S5.** BLAST analysis of amino acid sequences of uncharacterised proteins included within this study, identifying the characterised protein with the highest percentage amino acid sequence similarity for each uncharacterised protein.

| Accession Number of Uncharacterised Protein | Analysis Present In | Accession Number of Similar Protein | Similar Protein | Species | E-Value | Similarity (%) |
|---|---|---|---|---|---|---|
| F6PKE1 | HKJC Macroscopic OA Correlation<br>DAPC Linear Discriminant 2 | A0A340XC13 | Inhibitor of carbonic anhydrase-like isoform X1 | *Lipotes vexillifer* | 0 | 77.2 |
| F6S2I3 | Biobank Lasso Model | A0A1U7U041 | Glutathione S-transferase | *Tarsius syrichta* | 4.60 $e^{-144}$ | 88.5 |
| F6SP11 | Biobank Macroscopic OA Correlation | A0A0A1E4I0 | Immunoglobulin lambda light chain variable region | *Equus caballus* | 8.10 $e^{-60}$ | 100.0 |
| F6TED1 | Native (Macroscopic OA)<br>Biobank Lasso Model | A0A091E338 | Immunoglobulin kappa chain V-III region MOPC 63 | *Fukomys damarensis* | 2.20 $e^{-44}$ | 70.1 |
| F6ZR63 | Biobank Macroscopic OA Correlation<br>DAPC Linear Discriminant 1<br>Biobank Lasso Model | A0A383YWT8 | Complement factor H-like isoform X1 | *Balaenoptera acutorostrata scammoni* | 0 | 68.6 |
| F7APS1 | HKJC Macroscopic OA Correlation | Q862Z5 | Cystatin-B | *Macaca fuscata fuscata* | 2.30 $e^{-57}$ | 87.8 |
| F7BM69 | HKJC Macroscopic OA Correlation | A0A337SQC1 | Immunoglobulin kappa variable 4-1 | *Felis catus* | 1.50 $e^{-56}$ | 77.4 |
| F7DXM5 | HKJC Macroscopic OA Correlation | B5BV10 | Alpha-1-antitrypsin | *Equus caballus* | 0 | 98.1 |
| H9GZQ9 | Biobank Macroscopic OA Correlation<br>Biobank Lasso Model | Q95M34 | Immunoglobulin gamma 1 heavy chain constant region | *Equus caballus* | 0 | 99.7 |
| H9GZS6 | Native (Macroscopic OA)<br>Biobank Macroscopic OA Correlation | Q95M34 | Immunoglobulin gamma 1 heavy chain constant region | *Equus caballus* | 7.40 $e^{-173}$ | 70.1 |
| H9GZU9 | Native (Macroscopic OA)<br>HKJC Macroscopic OA Correlation<br>Biobank Macroscopic OA Correlation | Q95M34 | Immunoglobulin gamma 1 heavy chain constant region | *Equus caballus* | 1.10 $e^{-159}$ | 67.9 |
| H9GZV0 | Biobank Macroscopic OA Correlation | L5JR68 | Immunoglobulin epsilon chain C region | *Pteropus alecto* | 0 | 65.0 |

*Abbreviations: DAPC, Discriminant analysis of principal components; HKJC, Hong Kong Jockey Club; OA, Osteoarthritis*

**Table S6.** Macroscopic osteoarthritis scoring of distal metacarpal III for the University of Liverpool equine biobank.

| | Scorer 1 | | | | Scorer 2 | | | | Scorer 3 | | | | | |
|---|---|---|---|---|---|---|---|---|---|---|---|---|---|---|
| **Horse** | **Wear Lines (0-3)** | **Erosions (0-3)** | **Palmar Arthrosis (0-3)** | **Total** | **Wear Lines (0-3)** | **Erosions (0-3)** | **Palmar Arthrosis (0-3)** | **Total** | **Wear Lines (0-3)** | **Erosions (0-3)** | **Palmar Arthrosis (0-3)** | **Total** | **Average TOTAL** | **Macroscopic OA Grade** |
| **1** | Not Scored | | | | Not Scored | | | | | | | | | |
| **2** | 0 | 2 | 0 | **2** | 1 | 2 | 0 | **3** | | | | | 3 | **3** |
| **3** | 0 | 0 | 0 | **0** | 0 | 0 | 0 | **0** | | | | | 0 | **0** |
| **4** | Not Scored | | | | Not Scored | | | | | | | | | |
| **5** | Not Scored | | | | Not Scored | | | | | | | | | |
| **6** | Not Scored | | | | Not Scored | | | | | | | | | |
| **7** | 0 | 1 | 0 | **1** | 0 | 0 | 0 | **0** | | | | | 1 | **1** |
| **8** | 1 | 0 | 2 | **3** | 2 | 1 | 1 | **4** | | | | | 4 | **3** |
| **9** | 0 | 1 | 0 | **1** | 0 | 0 | 0 | **0** | | | | | 1 | **1** |
| **10** | 0 | 0 | 0 | **0** | 0 | 1 | 1 | **2** | 0 | 1 | 0 | **1** | 2 | **2** |
| **11** | 0 | 1 | 0 | **1** | 1 | 1 | 0 | **2** | | | | | 2 | **2** |
| **12** | 0 | 0 | 1 | **1** | 1 | 1 | 0 | **2** | | | | | 2 | **2** |
| **13** | 1 | 2 | 1 | **4** | 1 | 1 | 3 | **5** | | | | | 5 | **3** |
| **14** | 0 | 0 | 0 | **0** | 1 | 0 | 1 | **2** | 1 | 0 | 0 | **1** | 2 | **2** |
| **15** | Not Scored | | | | Not Scored | | | | | | | | | |
| **16** | 0 | 0 | 1 | **1** | 1 | 0 | 2 | **3** | 1 | 0 | 0 | **1** | 1 | **1** |
| **17** | Not Scored | | | | Not Scored | | | | | | | | | |
| **18** | 0 | 0 | 0 | **0** | 0 | 0 | 0 | **0** | | | | | 0 | **0** |
| **19** | 0 | 2 | 2 | **4** | 0 | 0 | 0 | **0** | 0 | 0 | 0 | **0** | 0 | **0** |
| **20** | 0 | 0 | 0 | **0** | 0 | 0 | 0 | **0** | | | | | 0 | **0** |
| **21** | 0 | 2 | 0 | **2** | 1 | 1 | 0 | **2** | | | | | 2 | **2** |
| **22** | 0 | 0 | 0 | **0** | 0 | 0 | 0 | **0** | | | | | 0 | **0** |
| **23** | 0 | 1 | 0 | **1** | 0 | 1 | 0 | **1** | | | | | 1 | **1** |
| **24** | 0 | 1 | 0 | **1** | 0 | 1 | 0 | **1** | | | | | 1 | **1** |
| **25** | 0 | 1 | 0 | **1** | 0 | 1 | 0 | **1** | | | | | 1 | **1** |
| **26** | 0 | 1 | 0 | **1** | 0 | 0 | 1 | **1** | | | | | 1 | **1** |
| **27** | 1 | 2 | 0 | **3** | 0 | 1 | 0 | **1** | 0 | 1 | 0 | **1** | 1 | **1** |
| **28** | 2 | 0 | 0 | **2** | 1 | 0 | 0 | **1** | | | | | 2 | **2** |
| **29** | 0 | 1 | 1 | **2** | 0 | 2 | 0 | **2** | | | | | 2 | **2** |
| **30** | 2 | 2 | 0 | **4** | 3 | 2 | 1 | **6** | 3 | 2 | 1 | **6** | 6 | **3** |

| | | | | | | | | | | | | | | |
|---|---|---|---|---|---|---|---|---|---|---|---|---|---|---|
| **31** | 0 | 2 | 0 | **2** | 0 | 2 | 0 | **2** | | | | | 2 | **2** |
| **32** | 3 | 1 | 0 | **4** | 2 | 1 | 0 | **3** | | | | | 4 | **3** |
| **33** | 1 | 1 | 0 | **2** | 0 | 0 | 1 | **1** | | | | | 2 | **2** |
| **34** | 1 | 1 | 0 | **2** | 0 | 1 | 0 | **1** | | | | | 2 | **2** |
| **35** | 2 | 2 | 0 | **4** | 3 | 2 | 2 | **7** | 2 | 1 | 2 | **5** | 5 | **3** |
| **36** | 0 | 0 | 1 | **1** | 0 | 0 | 1 | **1** | | | | | 1 | **1** |
| **37** | 0 | 0 | 0 | **0** | 0 | 0 | 1 | **1** | | | | | 1 | **1** |
| **38** | 0 | 1 | 0 | **1** | 0 | 2 | 0 | **2** | | | | | 2 | **2** |
| **39** | 0 | 2 | 1 | **3** | 1 | 2 | 1 | **4** | | | | | 4 | **3** |
| **40** | 0 | 1 | 0 | **1** | 0 | 2 | 0 | **2** | | | | | 2 | **2** |
| **41** | 0 | 2 | 0 | **2** | 0 | 1 | 0 | **1** | | | | | 2 | **2** |
| **42** | Not Scored | | | | Not Scored | | | | | | | | | |
| **43** | 0 | 1 | 0 | **1** | 0 | 0 | 1 | **1** | | | | | 1 | **1** |
| **44** | 0 | 0 | 0 | **0** | 0 | 1 | 0 | **1** | | | | | 1 | **1** |
| **45** | 1 | 0 | 0 | **1** | 0 | 0 | 0 | **0** | | | | | 1 | **1** |
| **46** | 0 | 0 | 0 | **0** | 0 | 0 | 0 | **0** | | | | | 0 | **0** |
| **47** | 0 | 0 | 1 | **1** | 1 | 2 | 0 | **3** | 0 | 0 | 1 | **1** | 1 | **1** |
| **48** | 2 | 2 | 0 | **4** | 2 | 3 | 0 | **5** | | | | | 5 | **3** |
| **49** | 1 | 2 | 0 | **3** | 0 | 2 | 0 | **2** | | | | | 3 | **3** |
| **50** | 0 | 0 | 0 | **0** | 0 | 1 | 0 | **1** | | | | | 1 | **1** |
| **51** | 1 | 2 | 0 | **3** | 0 | 2 | 0 | **2** | | | | | 3 | **3** |
| **52** | 0 | 0 | 0 | **0** | 0 | 0 | 0 | **0** | | | | | 0 | **0** |
| **53** | 0 | 2 | 0 | **2** | 0 | 1 | 0 | **1** | | | | | 2 | **2** |
| **54** | 0 | 0 | 0 | **0** | 0 | 0 | 0 | **0** | | | | | 0 | **0** |
| **55** | 0 | 0 | 0 | **0** | 0 | 1 | 0 | **1** | | | | | 1 | **1** |
| **56** | 1 | 0 | 0 | **1** | 0 | 1 | 0 | **1** | | | | | 1 | **1** |
| **57** | 0 | 0 | 0 | **0** | 0 | 0 | 0 | **0** | | | | | 0 | **0** |
| **58** | 0 | 0 | 1 | **1** | 0 | 2 | 0 | **2** | | | | | 2 | **2** |
| **59** | 0 | 0 | 0 | **0** | 0 | 1 | 0 | **1** | | | | | 1 | **1** |
| **60** | 0 | 1 | 0 | **1** | 0 | 1 | 0 | **1** | | | | | 1 | **1** |
| **61** | 0 | 0 | 0 | **0** | 0 | 1 | 0 | **1** | | | | | 1 | **1** |
| **62** | 0 | 0 | 0 | **0** | 0 | 1 | 0 | **1** | | | | | 1 | **1** |
| **63** | 0 | 0 | 0 | **0** | 0 | 0 | 0 | **0** | | | | | 0 | **0** |
| **64** | 0 | 0 | 0 | **0** | 0 | 1 | 0 | **1** | | | | | 1 | **1** |
| **65** | 0 | 2 | 0 | **2** | 0 | 1 | 0 | **1** | | | | | 2 | **2** |
| **66** | 0 | 1 | 0 | **1** | 0 | 1 | 0 | **1** | | | | | 1 | **1** |
| **67** | 0 | 0 | 0 | **0** | 0 | 0 | 0 | **0** | | | | | 0 | **0** |
| **68** | 1 | 2 | 0 | **3** | 1 | 1 | 1 | **3** | | | | | 3 | **3** |

| | | | | | | | | | | | | | | |
|---|---|---|---|---|---|---|---|---|---|---|---|---|---|---|
| **69** | 1 | 0 | 0 | **1** | 0 | 1 | 0 | **1** | | | | | 1 | **1** |
| **70** | 0 | 1 | 0 | **1** | 0 | 2 | 0 | **2** | | | | | 2 | **2** |
| **71** | 3 | 0 | 0 | **3** | 3 | 0 | 0 | **3** | | | | | 3 | **3** |
| **72** | 0 | 0 | 2 | **2** | 1 | 1 | 2 | **4** | 0 | 0 | 2 | **2** | 2 | **2** |
| **73** | 0 | 0 | 0 | **0** | 0 | 1 | 0 | **1** | | | | | 1 | **1** |
| **132** | 0 | 0 | 2 | **2** | 0 | 0 | 0 | **0** | 0 | 0 | 0 | **0** | 0 | **0** |
| **133** | 0 | 0 | 0 | **0** | 0 | 0 | 0 | **0** | | | | | 0 | **0** |
| **134** | 0 | 2 | 0 | **2** | 0 | 1 | 0 | **1** | | | | | 2 | **2** |
| **135** | 0 | 2 | 1 | **3** | 0 | 1 | 0 | **1** | 0 | 1 | 0 | **1** | 1 | **1** |
| **136** | 1 | 0 | 1 | **2** | 1 | 1 | 0 | **2** | | | | | 2 | **2** |
| **137** | 0 | 0 | 1 | **1** | 1 | 2 | 1 | **4** | 1 | 2 | 0 | **3** | 4 | **3** |
| **138** | 0 | 0 | 0 | **0** | 0 | 0 | 0 | **0** | | | | | 0 | **0** |
| **139** | 3 | 0 | 1 | **4** | 3 | 0 | 0 | **3** | | | | | 4 | **3** |
| **140** | 1 | 1 | 0 | **2** | 0 | 1 | 0 | **1** | | | | | 2 | **2** |
| **141** | 0 | 0 | 1 | **1** | 0 | 0 | 0 | **0** | | | | | 1 | **1** |

**Table S7.** Microscopic osteoarthritis scoring of distal metacarpal III for the University of Liverpool equine biobank.

| | Scorer 1 | | | | | | Scorer 2 | | | | | | Scorer 3 | | | | | | | |
|---|---|---|---|---|---|---|---|---|---|---|---|---|---|---|---|---|---|---|---|---|
| Horse | Fissuring (0-4) | Focal Cell Loss (0-4) | Chond. Nec. (0-4) | Chon. Form. (0-4) | Saff O Stain (0-4) | Total | Fissuring (0-4) | Focal Cell Loss (0-4) | Chond. Nec. (0-4) | Chon. Form. (0-4) | Saff O Stain (0-4) | Total | Fissuring (0-4) | Focal Cell Loss (0-4) | Chond. Nec. (0-4) | Chon. Form. (0-4) | Saff O Stain (0-4) | Total | Average TOTAL | Microscopic OA Grade |
| **1** | 0 | 1 | 1 | 0 | 0 | **2** | 1 | 1 | 0 | 0 | 0 | **2** | | | | | | | **2** | **1** |
| **2** | 2 | 1 | 0 | 1 | 0 | **4** | 2 | 0 | 1 | 1 | 0 | **4** | | | | | | | **4** | **2** |
| **3** | 0 | 1 | 0 | 0 | 0 | **1** | 0 | 0 | 0 | 0 | 0 | **0** | | | | | | | **1** | **0** |
| **4** | 0 | 0 | 1 | 0 | 0 | **1** | 0 | 1 | 1 | 0 | 0 | **2** | | | | | | | **2** | **1** |
| **5** | 1 | 1 | 0 | 0 | 0 | **2** | 1 | 1 | 0 | 1 | 0 | **3** | | | | | | | **3** | **1** |
| **6** | 0 | 0 | 0 | 0 | 0 | **0** | 0 | 2 | 1 | 0 | 0 | **3** | 0 | 0 | 0 | 0 | 0 | **0** | **0** | **0** |
| **7** | 1 | 1 | 0 | 0 | 0 | **2** | 1 | 0 | 1 | 0 | 1 | **3** | | | | | | | **3** | **1** |
| **8** | 1 | 0 | 1 | 1 | 0 | **3** | 2 | 1 | 1 | 1 | 0 | **5** | 2 | 1 | 0 | 1 | 0 | **4** | **5** | **2** |
| **9** | 1 | 0 | 0 | 0 | 1 | **2** | 0 | 2 | 2 | 0 | 1 | **5** | 0 | 1 | 0 | 0 | 0 | **1** | **2** | **1** |
| **10** | 0 | 0 | 0 | 1 | 0 | **1** | 1 | 1 | 0 | 0 | 0 | **2** | | | | | | | **2** | **1** |
| **11** | 1 | 2 | 0 | 0 | 0 | **3** | 1 | 0 | 0 | 0 | 0 | **1** | 1 | 0 | 0 | 0 | 0 | **1** | **1** | **0** |
| **12** | 2 | 2 | 1 | 0 | 0 | **5** | 2 | 1 | 2 | 0 | 0 | **5** | | | | | | | **5** | **2** |
| **13** | 1 | 1 | 0 | 0 | 0 | **2** | 1 | 1 | 0 | 0 | 0 | **2** | | | | | | | **2** | **1** |
| **14** | 1 | 1 | 1 | 2 | 0 | **5** | 1 | 3 | 0 | 1 | 0 | **5** | | | | | | | **5** | **2** |
| **15** | 1 | 1 | 0 | 0 | 0 | **2** | 1 | 1 | 0 | 0 | 0 | **2** | | | | | | | **2** | **1** |
| **16** | 0 | 2 | 0 | 1 | 0 | **3** | 0 | 1 | 0 | 1 | 0 | **2** | | | | | | | **3** | **1** |
| **17** | Not Scored | | | | | | Not Scored | | | | | | | | | | | | | |
| **18** | 0 | 0 | 0 | 0 | 0 | **0** | 0 | 1 | 0 | 0 | 0 | **1** | 0 | 1 | 0 | 0 | 0 | **1** | **1** | **0** |
| **19** | 1 | 2 | 0 | 1 | 0 | **4** | 0 | 1 | 0 | 0 | 0 | **1** | 1 | 0 | 0 | 0 | 0 | **1** | **1** | **0** |
| **20** | 0 | 2 | 0 | 0 | 0 | **2** | 0 | 2 | 0 | 0 | 0 | **2** | | | | | | | **2** | **1** |
| **21** | 2 | 0 | 1 | 2 | 1 | **6** | 3 | 1 | 1 | 1 | 2 | **8** | 1 | 3 | 0 | 1 | 2 | **7** | **8** | **2** |
| **22** | 0 | 1 | 0 | 0 | 0 | **1** | 0 | 0 | 1 | 0 | 0 | **1** | | | | | | | **1** | **0** |
| **23** | 1 | 0 | 2 | 0 | 0 | **3** | 0 | 0 | 0 | 0 | 0 | **0** | 2 | 1 | 0 | 0 | 0 | **3** | **3** | **1** |
| **24** | 0 | 0 | 0 | 1 | 0 | **1** | 1 | 0 | 0 | 1 | 0 | **2** | | | | | | | **2** | **1** |
| **25** | 0 | 1 | 1 | 0 | 0 | **2** | 0 | 1 | 1 | 1 | 0 | **3** | | | | | | | **3** | **1** |
| **26** | 0 | 1 | 0 | 0 | 0 | **1** | 0 | 1 | 1 | 0 | 0 | **2** | | | | | | | **2** | **1** |
| **27** | 0 | 0 | 0 | 0 | 0 | **0** | 0 | 0 | 0 | 0 | 0 | **0** | | | | | | | **0** | **0** |
| **28** | 1 | 0 | 0 | 1 | 0 | **2** | 1 | 3 | 0 | 0 | 0 | **4** | 1 | 2 | 1 | 0 | 0 | **4** | **4** | **2** |
| **29** | 1 | 1 | 0 | 0 | 0 | **2** | 0 | 0 | 0 | 1 | 0 | **1** | | | | | | | **2** | **1** |
| **30** | 2 | 1 | 1 | 1 | 0 | **5** | 1 | 0 | 0 | 2 | 0 | **3** | 0 | 1 | 0 | 3 | 0 | **4** | **5** | **2** |

| | | | | | | | | | | | | | | | | | | | | |
|---|---|---|---|---|---|---|---|---|---|---|---|---|---|---|---|---|---|---|---|---|
| **31** | 2 | 1 | 1 | 1 | 4 | **9** | 1 | 0 | 1 | 1 | 0 | **3** | 1 | 2 | 0 | 0 | 0 | **3** | **3** | **1** |
| **32** | 0 | 1 | 1 | 1 | 0 | **3** | 1 | 0 | 0 | 2 | 0 | **3** | | | | | | | **3** | **1** |
| **33** | 2 | 2 | 1 | 1 | 1 | **7** | 2 | 1 | 1 | 1 | 0 | **5** | 2 | 1 | 2 | 1 | 0 | **6** | **7** | **2** |
| **34** | 0 | 1 | 2 | 1 | 0 | **4** | 0 | 2 | 1 | 1 | 0 | **4** | | | | | | | **4** | **2** |
| **35** | 1 | 0 | 1 | 2 | 1 | **5** | 1 | 1 | 1 | 2 | 1 | **6** | | | | | | | **6** | **2** |
| **36** | 1 | 1 | 0 | 1 | 0 | **3** | 1 | 0 | 0 | 0 | 1 | **2** | | | | | | | **3** | **1** |
| **37** | 0 | 1 | 1 | 1 | 1 | **4** | 0 | 2 | 1 | 0 | 0 | **3** | | | | | | | **4** | **2** |
| **38** | 1 | 0 | 1 | 1 | 0 | **3** | 1 | 1 | 2 | 1 | 0 | **5** | 2 | 1 | 2 | 1 | 0 | **6** | **6** | **2** |
| **39** | 0 | 1 | 0 | 1 | 2 | **4** | 0 | 0 | 1 | 0 | 1 | **2** | 0 | 1 | 0 | 0 | 2 | **3** | **4** | **2** |
| **40** | 1 | 0 | 1 | 2 | 2 | **6** | 1 | 0 | 0 | 1 | 1 | **3** | 2 | 0 | 1 | 2 | 2 | **7** | **7** | **2** |
| **41** | 2 | 1 | 1 | 2 | 0 | **6** | 1 | 2 | 2 | 3 | 0 | **8** | 2 | 2 | 0 | 1 | 0 | **5** | **6** | **2** |
| **42** | 0 | 2 | 1 | 0 | 1 | **4** | 0 | 1 | 1 | 0 | 0 | **2** | 0 | 2 | 0 | 1 | 1 | **4** | **4** | **2** |
| **43** | 1 | 0 | 1 | 1 | 0 | **3** | 1 | 0 | 1 | 0 | 0 | **2** | | | | | | | **3** | **1** |
| **44** | 0 | 1 | 1 | 2 | 1 | **5** | 1 | 0 | 1 | 1 | 1 | **4** | | | | | | | **5** | **2** |
| **45** | 0 | 1 | 1 | 0 | 0 | **2** | 0 | 1 | 1 | 0 | 0 | **2** | | | | | | | **2** | **1** |
| **46** | 0 | 0 | 0 | 1 | 0 | **1** | 0 | 1 | 1 | 0 | 0 | **2** | | | | | | | **2** | **1** |
| **47** | 0 | 1 | 0 | 1 | 1 | **3** | 1 | 0 | 0 | 0 | 1 | **2** | | | | | | | **3** | **1** |
| **48** | 0 | 1 | 1 | 0 | 0 | **2** | 0 | 1 | 1 | 2 | 0 | **4** | 1 | 1 | 0 | 0 | 0 | **2** | **2** | **1** |
| **49** | 1 | 1 | 0 | 1 | 1 | **4** | 0 | 0 | 0 | 1 | 0 | **1** | 0 | 1 | 1 | 1 | 1 | **4** | **4** | **2** |
| **50** | 1 | 1 | 2 | 0 | 0 | **4** | 2 | 0 | 1 | 0 | 0 | **3** | | | | | | | **4** | **2** |
| **51** | 0 | 1 | 1 | 0 | 1 | **3** | 1 | 0 | 1 | 2 | 1 | **5** | 0 | 0 | 0 | 2 | 0 | **2** | **3** | **1** |
| **52** | Not Scored | | | | | | Not Scored | | | | | | | | | | | | | |
| **53** | 1 | 0 | 0 | 2 | 1 | **4** | 2 | 0 | 0 | 1 | 0 | **3** | | | | | | | **4** | **2** |
| **54** | 0 | 1 | 1 | 0 | 0 | **2** | 0 | 2 | 0 | 0 | 0 | **2** | | | | | | | **2** | **1** |
| **55** | 0 | 1 | 1 | 0 | 1 | **3** | 0 | 0 | 0 | 0 | 0 | **0** | 0 | 1 | 0 | 0 | 1 | **2** | **3** | **1** |
| **56** | 1 | 1 | 0 | 0 | 2 | **4** | 0 | 1 | 0 | 0 | 1 | **2** | 1 | 1 | 0 | 0 | 2 | **4** | **4** | **2** |
| **57** | 0 | 1 | 0 | 0 | 1 | **2** | 0 | 2 | 1 | 0 | 1 | **4** | 1 | 1 | 0 | 0 | 0 | **2** | **2** | **1** |
| **58** | 0 | 1 | 0 | 2 | 0 | **3** | 0 | 1 | 2 | 0 | 0 | **3** | | | | | | | **3** | **1** |
| **59** | 0 | 0 | 0 | 0 | 0 | **0** | 0 | 1 | 0 | 1 | 1 | **3** | 1 | 1 | 0 | 1 | 1 | **4** | **4** | **2** |
| **60** | 1 | 0 | 0 | 1 | 0 | **2** | 2 | 0 | 0 | 2 | 0 | **4** | 0 | 2 | 0 | 2 | 0 | **4** | **4** | **2** |
| **61** | 0 | 1 | 1 | 1 | 0 | **3** | 0 | 0 | 1 | 0 | 2 | **3** | | | | | | | **3** | **1** |
| **62** | 0 | 1 | 0 | 0 | 1 | **2** | 1 | 1 | 1 | 0 | 1 | **4** | 0 | 1 | 0 | 1 | 0 | **2** | **2** | **1** |
| **63** | 0 | 2 | 1 | 1 | 0 | **4** | 0 | 1 | 1 | 0 | 2 | **4** | | | | | | | **4** | **2** |
| **64** | 1 | 1 | 0 | 2 | 1 | **5** | 0 | 1 | 0 | 1 | 0 | **2** | 1 | 1 | 0 | 0 | 0 | **2** | **2** | **1** |
| **65** | 1 | 2 | 0 | 1 | 0 | **4** | 0 | 1 | 0 | 1 | 0 | **2** | 0 | 1 | 0 | 2 | 0 | **3** | **4** | **2** |
| **66** | 1 | 2 | 1 | 0 | 1 | **5** | 1 | 0 | 0 | 1 | 0 | **2** | 0 | 2 | 0 | 0 | 0 | **2** | **2** | **1** |
| **67** | 0 | 1 | 1 | 1 | 0 | **3** | 0 | 0 | 0 | 1 | 1 | **2** | | | | | | | **3** | **1** |
| **68** | 1 | 1 | 0 | 0 | 0 | **2** | 2 | 0 | 0 | 0 | 0 | **2** | | | | | | | **2** | **1** |

| | | | | | | | | | | | | | | | | | | | | |
|---|---|---|---|---|---|---|---|---|---|---|---|---|---|---|---|---|---|---|---|---|
| **69** | 2 | 0 | 1 | 1 | 0 | **4** | 1 | 0 | 1 | 1 | 0 | **3** | | | | | | | **4** | **2** |
| **70** | 3 | 1 | 1 | 1 | 2 | **8** | 2 | 0 | 1 | 1 | 2 | **6** | 3 | 2 | 0 | 0 | 2 | **7** | **8** | **2** |
| **71** | 2 | 2 | 1 | 1 | 0 | **6** | 1 | 1 | 0 | 1 | 0 | **3** | 1 | 2 | 0 | 3 | 0 | **6** | **6** | **2** |
| **72** | Not Scored | | | | | | Not Scored | | | | | | | | | | | | | |
| **73** | 2 | 1 | 1 | 0 | 0 | **4** | 0 | 0 | 0 | 0 | 0 | **0** | 1 | 0 | 0 | 0 | 0 | **1** | **1** | **0** |
| **132** | Not Scored | | | | | | Not Scored | | | | | | | | | | | | | |
| **133** | Not Scored | | | | | | Not Scored | | | | | | | | | | | | | |
| **134** | Not Scored | | | | | | Not Scored | | | | | | | | | | | | | |
| **135** | Not Scored | | | | | | Not Scored | | | | | | | | | | | | | |
| **136** | Not Scored | | | | | | Not Scored | | | | | | | | | | | | | |
| **137** | Not Scored | | | | | | Not Scored | | | | | | | | | | | | | |
| **138** | Not Scored | | | | | | Not Scored | | | | | | | | | | | | | |
| **139** | Not Scored | | | | | | Not Scored | | | | | | | | | | | | | |
| **140** | Not Scored | | | | | | Not Scored | | | | | | | | | | | | | |
| **141** | Not Scored | | | | | | Not Scored | | | | | | | | | | | | | |

**Table S8.** Macroscopic osteoarthritis scoring of distal metacarpal III or metatarsal III for the Hong Kong Jockey Club sample set.

| | | Scorer 1 | | | | | Scorer 2 | | | | | Scorer 3 | | | | | | |
|---|---|---|---|---|---|---|---|---|---|---|---|---|---|---|---|---|---|---|
| Horse | Joint | POD (0-3) | Wear lines (0-2) | Cart. loss (0-3) | Linear fissures (0-3) | TOTAL | POD (0-3) | Wear lines (0-2) | Cart. loss (0-3) | Linear fissures (0-3) | TOTAL | POD (0-3) | Wear lines (0-2) | Cart. loss (0-3) | Linear fissures (0-3) | TOTAL | Average TOTAL | Macroscopic OA Grade |
| 74 | MCP | Not Scored | | | | | Not Scored | | | | | | | | | | | |
| 75 | MCP | 2 | 0 | 3 | 0 | 5 | 1 | 0 | 1 | 0 | 2 | 2 | 0 | 2 | 0 | 4 | 5 | **2** |
| 76 | MCP | 0 | 2 | 2 | 1 | 5 | 2 | 2 | 1 | 0 | 5 | | | | | | 5 | **2** |
| 77 | MCP | 3 | 2 | 3 | 1 | 9 | 3 | 2 | 3 | 0 | 8 | | | | | | 9 | **2** |
| 78 | MCP | 0 | 2 | 0 | 2 | 4 | 0 | 1 | 0 | 0 | 1 | 0 | 1 | 0 | 1 | 2 | 2 | **0** |
| 79 | MCP | 0 | 1 | 0 | 2 | 3 | 0 | 1 | 1 | 0 | 2 | | | | | | 3 | **1** |
| 80 | MCP | 3 | 2 | 1 | 0 | 6 | 3 | 2 | 1 | 0 | 6 | | | | | | 6 | **2** |
| 81 | MCP | Not Scored | | | | | Not Scored | | | | | | | | | | | |
| 82 | MCP | 2 | 1 | 0 | 0 | 3 | 2 | 1 | 1 | 0 | 4 | | | | | | 4 | **1** |
| 83 | MCP | 1 | 2 | 1 | 0 | 4 | 1 | 2 | 1 | 0 | 4 | | | | | | 4 | **1** |
| 84 | MCP | 0 | 0 | 3 | 1 | 4 | 0 | 0 | 1 | 1 | 2 | 0 | 1 | 1 | 1 | 3 | 4 | **1** |
| 85 | MCP | 0 | 2 | 3 | 0 | 5 | 0 | 2 | 1 | 0 | 3 | 0 | 2 | 1 | 0 | 3 | 3 | **1** |
| 86 | MCP | 2 | 2 | 3 | 0 | 7 | 2 | 1 | 1 | 0 | 4 | 1 | 1 | 1 | 0 | 3 | 4 | **1** |
| 87 | MCP | 3 | 2 | 3 | 0 | 8 | 3 | 2 | 1 | 0 | 6 | 2 | 1 | 2 | 0 | 5 | 6 | **2** |
| 88 | MCP | 1 | 0 | 0 | 0 | 1 | 1 | 0 | 0 | 2 | 3 | 1 | 1 | 0 | 1 | 3 | 3 | **1** |
| 89 | MCP | 3 | 2 | 3 | 0 | 8 | 3 | 2 | 1 | 0 | 6 | 3 | 2 | 3 | 0 | 8 | 8 | **2** |
| 90 | MCP | 1 | 0 | 3 | 0 | 4 | 1 | 0 | 1 | 0 | 2 | 1 | 1 | 1 | 0 | 3 | 4 | **1** |
| 91 | MCP | 1 | 1 | 2 | 0 | 4 | 1 | 1 | 1 | 0 | 3 | | | | | | 4 | **1** |
| 92 | MCP | 3 | 2 | 3 | 0 | 8 | 1 | 1 | 1 | 0 | 3 | | | | | | 6 | **2** |
| 93 | MCP | 2 | 0 | 3 | 0 | 5 | 2 | 0 | 2 | 1 | 5 | | | | | | 5 | **2** |
| 94 | MCP | 0 | 0 | 2 | 0 | 2 | 1 | 1 | 1 | 0 | 3 | | | | | | 3 | **1** |
| 95 | MCP | 2 | 1 | 3 | 3 | 9 | 1 | 1 | 1 | 3 | 6 | 2 | 1 | 2 | 1 | 6 | 6 | **2** |
| 96 | MCP | 2 | 0 | 3 | 0 | 5 | 2 | 0 | 1 | 1 | 4 | | | | | | 5 | **2** |
| 97 | MCP | 2 | 2 | 1 | 0 | 5 | 2 | 1 | 0 | 0 | 3 | 1 | 2 | 1 | 0 | 4 | 5 | **2** |
| 98 | MCP | 0 | 0 | 0 | 1 | 1 | 0 | 0 | 0 | 1 | 1 | | | | | | 1 | **0** |
| 99 | MCP | 0 | 0 | 0 | 0 | 0 | 0 | 1 | 0 | 1 | 2 | 0 | 0 | 0 | 0 | 0 | 0 | **0** |
| 100 | MCP | 0 | 1 | 0 | 2 | 3 | 1 | 1 | 0 | 1 | 3 | | | | | | 3 | **1** |
| 101 | MCP | 1 | 0 | 0 | 0 | 1 | 0 | 0 | 1 | 1 | 2 | | | | | | 2 | **0** |
| 102 | MCP | 1 | 1 | 2 | 0 | 4 | 1 | 1 | 2 | 1 | 5 | | | | | | 5 | **2** |
| 103 | MCP | 0 | 0 | 0 | 1 | 1 | 0 | 0 | 1 | 1 | 2 | | | | | | 2 | **0** |

| | | | | | | | | | | | | | | | | | | |
|---|---|---|---|---|---|---|---|---|---|---|---|---|---|---|---|---|---|---|
| 104 | MCP | Not Scored | | | | | Not Scored | | | | | | | | | | | |
| 105 | MTP | Not Scored | | | | | Not Scored | | | | | | | | | | | |
| 106 | MTP | 0 | 0 | 2 | 2 | 4 | 0 | 0 | 1 | 1 | 2 | 0 | 0 | 0 | 1 | 1 | 2 | **0** |
| 107 | MTP | Not Scored | | | | | Not Scored | | | | | | | | | | | |
| 108 | MTP | 0 | 0 | 1 | 2 | 3 | 1 | 0 | 1 | 1 | 3 | | | | | | 3 | **1** |
| 109 | MTP | 2 | 0 | 1 | 2 | 5 | 2 | 0 | 1 | 2 | 5 | | | | | | 5 | **2** |
| 110 | MTP | 3 | 2 | 3 | 0 | 8 | 0 | 0 | 0 | 1 | 1 | 0 | 1 | 1 | 0 | 2 | 2 | **0** |
| 111 | MTP | 0 | 2 | 2 | 0 | 4 | 1 | 1 | 0 | 1 | 3 | | | | | | 4 | **1** |
| 112 | MTP | Not Scored | | | | | Not Scored | | | | | | | | | | | |
| 113 | MTP | 0 | 1 | 3 | 1 | 5 | 1 | 1 | 0 | 1 | 3 | 0 | 1 | 0 | 2 | 3 | 3 | **1** |
| 114 | MTP | 1 | 0 | 1 | 0 | 2 | 1 | 0 | 1 | 1 | 3 | | | | | | 3 | **1** |
| 115 | MTP | 0 | 1 | 0 | 3 | 4 | 1 | 1 | 0 | 2 | 4 | | | | | | 4 | **1** |
| 116 | MTP | 0 | 0 | 0 | 3 | 3 | 0 | 0 | 0 | 2 | 2 | | | | | | 3 | **1** |
| 117 | MTP | 0 | 0 | 0 | 3 | 3 | 0 | 1 | 0 | 1 | 2 | | | | | | 3 | **1** |
| 118 | MTP | 0 | 0 | 0 | 2 | 2 | 1 | 0 | 0 | 2 | 3 | | | | | | 3 | **1** |
| 119 | MTP | 0 | 0 | 1 | 2 | 3 | 0 | 0 | 0 | 2 | 2 | | | | | | 3 | **1** |
| 120 | MTP | 1 | 1 | 0 | 0 | 2 | 0 | 1 | 0 | 2 | 3 | | | | | | 3 | **1** |
| 121 | MTP | 0 | 0 | 0 | 1 | 1 | 0 | 0 | 0 | 1 | 1 | | | | | | 1 | **0** |
| 122 | MTP | 0 | 0 | 0 | 3 | 3 | 0 | 0 | 0 | 1 | 1 | | | | | | 2 | **0** |
| 123 | MTP | 1 | 1 | 0 | 0 | 2 | 1 | 0 | 0 | 2 | 3 | | | | | | 3 | **1** |
| 124 | MTP | Not Scored | | | | | Not Scored | | | | | | | | | | | |
| 125 | MTP | 2 | 0 | 0 | 2 | 4 | 1 | 0 | 0 | 1 | 2 | | | | | | 3 | **1** |
| 126 | MTP | Not Scored | | | | | Not Scored | | | | | | | | | | | |
| 127 | MTP | 0 | 0 | 0 | 1 | 1 | 0 | 0 | 0 | 2 | 2 | | | | | | 2 | **0** |
| 128 | MTP | 0 | 0 | 0 | 2 | 2 | 0 | 0 | 0 | 2 | 2 | | | | | | 2 | **0** |
| 129 | MTP | 1 | 0 | 0 | 1 | 2 | 1 | 0 | 0 | 2 | 3 | | | | | | 3 | **1** |
| 130 | MTP | 0 | 0 | 0 | 2 | 2 | 0 | 0 | 0 | 1 | 1 | | | | | | 2 | **0** |
| 131 | MCP | 0 | 1 | 3 | 1 | 5 | 1 | 1 | 1 | 0 | 3 | | | | | | 4 | **1** |

**Table S9.** Microscopic osteoarthritis scoring of distal metacarpal III or metatarsal III for the Hong Kong Jockey Club sample set.

| Horse | Joint | Structure (0-10) | Cell Density (0-4) | Cell Cloning (0-4) | Staining (0-4) | Tidemark (0-3) | TOTAL | Microscopic OA Grade |
|---|---|---|---|---|---|---|---|---|
| 74 | MCP | Not Scored | | | | | | |
| 75 | MCP | 1 | 0 | 2 | 2 | 3 | 8 | **1** |
| 76 | MCP | 8 | 1 | 4 | 4 | 3 | 20 | **2** |
| 77 | MCP | 1 | 0 | 1 | 1 | 2 | 5 | **1** |
| 78 | MCP | 2 | 0 | 0 | 1 | 1 | 4 | **0** |
| 79 | MCP | 1 | 1 | 1 | 0 | 0 | 3 | **0** |
| 80 | MCP | Not Scored | | | | | | |
| 81 | MCP | Not Scored | | | | | | |
| 82 | MCP | 8 | 4 | 4 | 2 | 1 | 19 | **2** |
| 83 | MCP | 6 | 2 | 3 | 4 | 1 | 16 | **2** |
| 84 | MCP | 0 | 1 | 1 | 0 | 0 | 2 | **0** |
| 85 | MCP | 1 | 1 | 1 | 2 | 1 | 6 | **1** |
| 86 | MCP | 1 | 2 | 2 | 1 | 2 | 8 | **1** |
| 87 | MCP | 0 | 0 | 0 | 0 | 0 | 0 | **0** |
| 88 | MCP | 3 | 0 | 2 | 2 | 3 | 10 | **1** |
| 89 | MCP | 2 | 1 | 1 | 3 | 2 | 9 | **1** |
| 90 | MCP | 7 | 1 | 0 | 3 | 3 | 14 | **2** |
| 91 | MCP | 7 | 1 | 1 | 4 | 3 | 16 | **2** |
| 92 | MCP | 10 | 1 | 0 | 4 | 3 | 18 | **2** |
| 93 | MCP | 4 | 2 | 0 | 2 | 3 | 11 | **2** |
| 94 | MCP | 2 | 1 | 2 | 1 | 3 | 9 | **1** |
| 95 | MCP | 10 | 1 | 2 | 2 | 3 | 18 | **2** |
| 96 | MCP | Not Scored | | | | | | |
| 97 | MCP | 1 | 0 | 1 | 0 | 1 | 3 | **0** |
| 98 | MCP | 0 | 0 | 1 | 0 | 0 | 1 | **0** |
| 99 | MCP | 2 | 1 | 0 | 0 | 0 | 3 | **0** |
| 100 | MCP | Not Scored | | | | | | |
| 101 | MCP | 3 | 0 | 0 | 2 | 2 | 7 | **1** |
| 102 | MCP | 10 | 2 | 0 | 3 | 3 | 18 | **2** |
| 103 | MCP | 1 | 0 | 0 | 1 | 2 | 4 | **0** |
| 104 | MCP | 2 | 0 | 2 | 0 | 0 | 4 | **0** |
| 105 | MTP | Not Scored | | | | | | |
| 106 | MTP | 1 | 0 | 0 | 1 | 0 | 2 | **0** |
| 107 | MTP | 2 | 0 | 1 | 2 | 2 | 7 | **1** |
| 108 | MTP | 5 | 0 | 0 | 2 | 3 | 10 | **1** |
| 109 | MTP | 2 | 1 | 1 | 1 | 0 | 5 | **1** |
| 110 | MTP | 4 | 2 | 0 | 4 | 2 | 12 | **2** |
| 111 | MTP | 5 | 1 | 1 | 3 | 2 | 12 | **2** |
| 112 | MTP | Not Scored | | | | | | |
| 113 | MTP | 2 | 0 | 1 | 1 | 1 | 5 | **1** |
| 114 | MTP | 2 | 1 | 3 | 4 | 2 | 12 | **2** |
| 115 | MTP | 3 | 3 | 4 | 1 | 1 | 12 | **2** |
| 116 | MTP | 2 | 0 | 1 | 0 | 0 | 3 | **0** |
| 117 | MTP | 1 | 1 | 1 | 2 | 2 | 7 | **1** |
| 118 | MTP | 3 | 0 | 1 | 3 | 1 | 8 | **1** |
| 119 | MTP | 1 | 0 | 0 | 1 | 0 | 2 | **0** |
| 120 | MTP | 1 | 0 | 1 | 1 | 0 | 3 | **0** |

| | | | | | | | | |
|---|---|---|---|---|---|---|---|---|
| 121 | MTP | 1 | 1 | 1 | 2 | 3 | 8 | **1** |
| 122 | MTP | 1 | 1 | 1 | 0 | 0 | 3 | **0** |
| 123 | MTP | 1 | 0 | 0 | 3 | 3 | 7 | **1** |
| 124 | MTP | Not Scored | | | | | | |
| 125 | MTP | 2 | 1 | 1 | 2 | 0 | 6 | **1** |
| 126 | MTP | 2 | 0 | 0 | 1 | 2 | 5 | **1** |
| 127 | MTP | 2 | 1 | 1 | 2 | 2 | 8 | **1** |
| 128 | MTP | 1 | 0 | 1 | 1 | 3 | 6 | **1** |
| 129 | MTP | 1 | 0 | 0 | 1 | 2 | 4 | **0** |
| 130 | MTP | 1 | 1 | 1 | 1 | 0 | 4 | **0** |
| 131 | MCP | 0 | 1 | 1 | 1 | 0 | 3 | **0** |

**Table S10.** Synovitis scoring of distal metacarpal III or metatarsal III for the Hong Kong Jockey Club sample set.

| | | Lining Cell Layer (0-3) | | | | Resident Cells (0-3) | | | | Inflammatory infiltrate (0-3) | | | | | |
|---|---|---|---|---|---|---|---|---|---|---|---|---|---|---|---|
| Horse | Joint | 1 | 2 | 3 | Average | 1 | 2 | 3 | Average | 1 | 2 | 3 | Average | Total | Synovitis Grade |
| 74 | MCP | 1 | 1 | 0 | **0.7** | 2 | 2 | 0 | **1.3** | 2 | 2 | 1 | **1.7** | **4** | **1** |
| 75 | MCP | 1 | 1 | 1 | **1.0** | 1 | 2 | 1 | **1.3** | 0 | 0 | 1 | **0.3** | **3** | **1** |
| 76 | MCP | 2 | 2 | 1 | **1.7** | 1 | 1 | 1 | **1.0** | 1 | 2 | 1 | **1.3** | **4** | **1** |
| 77 | MCP | 2 | 1 | 3 | **2.0** | 2 | 2 | 1 | **1.7** | 1 | 1 | 1 | **1.0** | **5** | **2** |
| 78 | MCP | 2 | 1 | 1 | **1.3** | 1 | 1 | 1 | **1.0** | 1 | 1 | 1 | **1.0** | **3** | **1** |
| 79 | MCP | 1 | 2 | 2 | **1.7** | 1 | 1 | 1 | **1.0** | 1 | 1 | 1 | **1.0** | **4** | **1** |
| 80 | MCP | 1 | 2 | 2 | **1.7** | 1 | 1 | 1 | **1.0** | 1 | 1 | 1 | **1.0** | **4** | **1** |
| 81 | MCP | 0 | 1 | 1 | **0.7** | 1 | 2 | 2 | **1.7** | 0 | 3 | 3 | **2.0** | **4** | **1** |
| 82 | MCP | 1 | 1 | 3 | **1.7** | 1 | 2 | 2 | **1.7** | 2 | 2 | 3 | **2.3** | **6** | **2** |
| 83 | MCP | 2 | 1 | 2 | **1.7** | 1 | 1 | 1 | **1.0** | 1 | 1 | 1 | **1.0** | **4** | **1** |
| 84 | MCP | 1 | 1 | 2 | **1.3** | 1 | 1 | 1 | **1.0** | 1 | 1 | 1 | **1.0** | **3** | **1** |
| 85 | MCP | 2 | 2 | 2 | **2.0** | 2 | 1 | 1 | **1.3** | 1 | 1 | 1 | **1.0** | **4** | **1** |
| 86 | MCP | 2 | 2 | 1 | **1.7** | 1 | 1 | 1 | **1.0** | 1 | 1 | 1 | **1.0** | **4** | **1** |
| 87 | MCP | 2 | 3 | 1 | **2.0** | 3 | 2 | 3 | **2.7** | 0 | 1 | 0 | **0.3** | **5** | **2** |
| 88 | MCP | 2 | 2 | 2 | **2.0** | 1 | 1 | 1 | **1.0** | 2 | 1 | 1 | **1.3** | **4** | **1** |
| 89 | MCP | 2 | 3 | 2 | **2.3** | 2 | 2 | 2 | **2.0** | 2 | 2 | 2 | **2.0** | **6** | **2** |
| 90 | MCP | 3 | 2 | 3 | **2.7** | 3 | 3 | 3 | **3.0** | 2 | 2 | 2 | **2.0** | **8** | **2** |
| 91 | MCP | 1 | 2 | 2 | **1.7** | 1 | 2 | 1 | **1.3** | 1 | 1 | 2 | **1.3** | **4** | **1** |
| 92 | MCP | 2 | 3 | 3 | **2.7** | 1 | 1 | 2 | **1.3** | 1 | 1 | 2 | **1.3** | **5** | **2** |
| 93 | MCP | 3 | 2 | 2 | **2.3** | 2 | 1 | 3 | **2.0** | 1 | 2 | 1 | **1.3** | **6** | **2** |
| 94 | MCP | 0 | 1 | 1 | **0.7** | 1 | 2 | 1 | **1.3** | 1 | 1 | 2 | **1.3** | **3** | **1** |
| 95 | MCP | 2 | 2 | 3 | **2.3** | 3 | 2 | 3 | **2.7** | 2 | 2 | 2 | **2.0** | **7** | **2** |
| 96 | MCP | 1 | 1 | 3 | **1.7** | 2 | 2 | 2 | **2.0** | 1 | 1 | 1 | **1.0** | **5** | **2** |
| 97 | MCP | 1 | 2 | 3 | **2.0** | 1 | 1 | 1 | **1.0** | 1 | 1 | 1 | **1.0** | **4** | **1** |
| 98 | MCP | 2 | 1 | 1 | **1.3** | 1 | 1 | 1 | **1.0** | 1 | 1 | 1 | **1.0** | **3** | **1** |
| 99 | MCP | 1 | 1 | 1 | **1.0** | 1 | 1 | 1 | **1.0** | 1 | 1 | 1 | **1.0** | **3** | **1** |
| 100 | MCP | 0 | 1 | 1 | **0.7** | 1 | 1 | 1 | **1.0** | 0 | 1 | 1 | **0.7** | **2** | **1** |
| 101 | MCP | 0 | 0 | 0 | **0.0** | 1 | 1 | 1 | **1.0** | 1 | 1 | 1 | **1.0** | **2** | **1** |
| 102 | MCP | 3 | 3 | 3 | **3.0** | 2 | 2 | 3 | **2.3** | 2 | 2 | 1 | **1.7** | **7** | **2** |
| 103 | MCP | 1 | 0 | 2 | **1.0** | 1 | 1 | 1 | **1.0** | 2 | 2 | 1 | **1.7** | **4** | **1** |
| 104 | MCP | 0 | 1 | 2 | **1.0** | 0 | 1 | 1 | **0.7** | 1 | 1 | 1 | **1.0** | **3** | **1** |
| 105 | MTP | 1 | 2 | 2 | **1.7** | 1 | 3 | 3 | **2.3** | 2 | 2 | 2 | **2.0** | **6** | **2** |
| 106 | MTP | 2 | 1 | 2 | **1.7** | 1 | 1 | 1 | **1.0** | 1 | 1 | 1 | **1.0** | **4** | **1** |
| 107 | MTP | 1 | 2 | 1 | **1.3** | 1 | 1 | 1 | **1.0** | 1 | 1 | 1 | **1.0** | **3** | **1** |
| 108 | MTP | 1 | 1 | 0 | **0.7** | 0 | 1 | 0 | **0.3** | 1 | 1 | 1 | **1.0** | **2** | **1** |
| 109 | MTP | 2 | 2 | 3 | **2.3** | 1 | 1 | 1 | **1.0** | 1 | 1 | 1 | **1.0** | **4** | **1** |
| 110 | MTP | 3 | 1 | 0 | **1.3** | 1 | 1 | 0 | **0.7** | 1 | 2 | 1 | **1.3** | **3** | **1** |
| 111 | MTP | 2 | 2 | 0 | **1.3** | 1 | 1 | 1 | **1.0** | 1 | 1 | 1 | **1.0** | **3** | **1** |
| 112 | MTP | 2 | 2 | 1 | **1.7** | 1 | 1 | 1 | **1.0** | 2 | 1 | 2 | **1.7** | **4** | **1** |
| 113 | MTP | 2 | 3 | 1 | **2.0** | 1 | 2 | 0 | **1.0** | 1 | 3 | 0 | **1.3** | **4** | **1** |
| 114 | MTP | 1 | 1 | 1 | **1.0** | 1 | 1 | 1 | **1.0** | 1 | 1 | 1 | **1.0** | **3** | **1** |
| 115 | MTP | 0 | 0 | 1 | **0.3** | 0 | 0 | 0 | **0.0** | 1 | 1 | 1 | **1.0** | **1** | **0** |
| 116 | MTP | 2 | 3 | 3 | **2.7** | 1 | 1 | 1 | **1.0** | 1 | 1 | 1 | **1.0** | **5** | **2** |
| 117 | MTP | 1 | 1 | 2 | **1.3** | 0 | 1 | 1 | **0.7** | 1 | 1 | 1 | **1.0** | **3** | **1** |
| 118 | MTP | 3 | 2 | 2 | **2.3** | 2 | 1 | 2 | **1.7** | 1 | 1 | 1 | **1.0** | **5** | **2** |
| 119 | MTP | 2 | 2 | 2 | **2.0** | 2 | 3 | 1 | **2.0** | 2 | 2 | 2 | **2.0** | **6** | **2** |

| | | | | | | | | | | | | | | | |
|---|---|---|---|---|---|---|---|---|---|---|---|---|---|---|---|
| 120 | MTP | 1 | 1 | 1 | **1.0** | 1 | 1 | 1 | **1.0** | 1 | 1 | 1 | **1.0** | **3** | **1** |
| 121 | MTP | 0 | 2 | 2 | **1.3** | 1 | 1 | 3 | **1.7** | 2 | 2 | 3 | **2.3** | **5** | **2** |
| 122 | MTP | 2 | 1 | 3 | **2.0** | 1 | 1 | 1 | **1.0** | 1 | 1 | 2 | **1.3** | **4** | **1** |
| 123 | MTP | 2 | 2 | 1 | **1.7** | 1 | 1 | 1 | **1.0** | 2 | 1 | 1 | **1.3** | **4** | **1** |
| 124 | MTP | 2 | 2 | 2 | **2.0** | 1 | 2 | 1 | **1.3** | 1 | 1 | 1 | **1.0** | **4** | **1** |
| 125 | MTP | 2 | 2 | 2 | **2.0** | 1 | 1 | 0 | **0.7** | 1 | 1 | 1 | **1.0** | **4** | **1** |
| 126 | MTP | 1 | 2 | 1 | **1.3** | 1 | 1 | 1 | **1.0** | 1 | 1 | 1 | **1.0** | **3** | **1** |
| 127 | MTP | 0 | 2 | 2 | **1.3** | 1 | 2 | 2 | **1.7** | 1 | 2 | 2 | **1.7** | **5** | **2** |
| 128 | MTP | 1 | 3 | 3 | **2.3** | 1 | 1 | 2 | **1.3** | 1 | 3 | 3 | **2.3** | **6** | **2** |
| 129 | MTP | 2 | 2 | 1 | **1.7** | 2 | 2 | 1 | **1.7** | 1 | 2 | 1 | **1.3** | **5** | **2** |
| 130 | MTP | 3 | 3 | 2 | **2.7** | 1 | 1 | 1 | **1.0** | 1 | 1 | 1 | **1.0** | **5** | **2** |
| 131 | MCP | 1 | 3 | 2 | **2.0** | 1 | 1 | 1 | **1.0** | 1 | 1 | 1 | **1.0** | **4** | **1** |
